# Specialization of the photoreceptor transcriptome by *Srrm3*-dependent microexons is required for outer segment maintenance and vision

**DOI:** 10.1101/2021.09.08.459463

**Authors:** Ludovica Ciampi, Federica Mantica, Laura Lopez-Blanch, Cristina Rodríguez-Marin, Damiano Cianferoni, Jingjing Zang, Jon Permanyer, Senda Jiménez-Delgado, Sophie Bonnal, Samuel Miravet-Verde, Verena Ruprecht, Stephan C.F. Neuhauss, Sandro Banfi, Sabrina Carrella, Luis Serrano, Sarah A. Head, Manuel Irimia

**Affiliations:** Centre for Genomic Regulation (CRG), The Barcelona Institute of Science and Technology, Barcelona, Spain; Universitat Pompeu Fabra (UPF), Barcelona, Spain; ICREA, Pg. Lluís Companys 23, Barcelona 08010, Spain; Department of Molecular Life Sciences, University of Zurich, Winterthurerstrasse 190, CH-8057 Zurich Switzerland; Medical Genetics, Department of Precision Medicine, University of Campania “L. Vanvitelli”, Naples, Italy; Telethon Institute of Genetics and Medicine (TIGEM), Pozzuoli (NA), Italy

## Abstract

Retinal photoreceptors differ in their transcriptomic profiles from other neuronal subtypes, likely as a reflection of their unique cellular morphology and function in the detection of light thorough the ciliary outer segment. We discovered a new layer of this molecular specialization by revealing that the vertebrate retina expresses the largest number of tissue-enriched microexons of all tissue types. A subset of these microexons is included exclusively in photoreceptor transcripts, particularly in genes involved in cilia biogenesis and in vesicle-mediated transport. This microexon program is regulated by *Srrm3*, a paralog of the neural microexon regulator *Srrm4*. Despite both proteins positively regulate retina microexons *in vitro*, only *Srrm3* is highly expressed in mature photoreceptors and its deletion in zebrafish results in widespread downregulation of microexon inclusion, severe photoreceptor alterations and blindness. These results shed light into photoreceptor’s transcriptomic specialization and functionality, uncovering new cell type-specific roles for *Srrm3* and microexons with implication for retinal diseases.

## INTRODUCTION

Impaired vision is a highly heterogeneous condition affecting millions of individuals worldwide. The key feature that accounts for most visual disabilities is the primary or secondary loss of photoreceptor (PR) cells, arising from a number of different genetic and environmental causes^1^. PRs, comprising rods and cones, have a unique ciliary structure named the Outer Segment (OS) that physically supports the phototransduction cascade, the process by which incoming light is converted into electrical signals that the brain can process^2^. This ciliary function contrasts with those of other cell types, which are generally involved in sensing vibration, liquid circulation, hormones, chemicals or temperature, and it is reflected by unique morphological characteristics. OSs are filled with densely packed and organized membranous disks that undergo rapid and extensive renewal to guarantee the proper supply of proteins such as rhodopsin, cone opsins and other visual pigments^3^. The delivery of those components from the Golgi to the ciliary tip and back is sustained by a high-capacity vesicle-mediated transport system, carried along the ciliary axoneme^4^. Consequently, multiple genes whose mutations are known to cause PR degeneration leading to syndromic or non-syndromic retinal ciliopathies encode for proteins involved in OS vesicular transport^4,5^.

Alternative splicing is a pre-translational mechanism used by specialized cell types to remodel their transcriptomes to produce protein isoforms necessary for their activity. Indeed, the high degree of functional specialization of PR cells is mirrored by the unusually large fraction of genes that undergo retina-specific alternative splicing, thereby creating unique isoforms in PRs essential for different cellular properties such as OS biogenesis^6^. The unique pattern of alternative splicing in PRs has been described and is known to be at least in part driven by the action of an RNA binding protein called Musashi 1 (*MSI1*)^7^. Recently, *MSI1* expression, combined with downregulation of the splicing factors *PTBP1* and *PCBP2*, has been linked to the activation of PR-specific exons^8^. Considering that aberrant splicing due to mutations in splicing factors and in retina-specific exons have all been linked to several retinal diseases (e.g. Retinitis Pigmentosa^9^), a complete characterization of the retinal alternative splicing programs is of great benefit for precision medicine.

Microexons are very short exons, ranging from 3 to 27 nucleotides, that are evolutionarily conserved among vertebrates and enriched in neurons^10^. These exons can encode as little as one amino acid and are often located on the surface of proteins where they can modulate protein-protein interactions^10-13^. The neuronal-specific Ser/Arg repetitive matrix protein 4 (*SRRM4*) controls the inclusion of most neuronal microexons annotated so far^10,14^. Depletion of *SRRM4* leads to neurodevelopmental defects in cell cultures as well as in *in vivo* models^15-18^. Recently, a neurally-enriched vertebrate-specific paralog of *SRRM4*, called *SRRM3*, has been reported^19^. *Srrm3* gene-trapped (Srrm3^gt/gt^) mice show reduced body size and lifespan, as well as tremors and ataxia. These mice also exhibit neuronal splicing defects, which become more severe when *SRRM4* expression levels are low^19^. *SRRM4* and *SRRM3* regulate a highly overlapping set of alternatively spliced small exons *in vitro*, and they share a 39 amino acid domain (enhancer of microexons, or eMIC) at their C-termini that is necessary and sufficient for the inclusion of neural microexons^14^. However, the roles of *SRRM4* and *SRRM3* in PRs remain unexplored.

Here, we find that the human retina expresses the largest program of tissue-enriched microexons among all tissue types, with a subset of those included only in PRs. Retina-enriched microexons (hereafter, RetMICs) are enriched in genes involved in cilia biogenesis and vesicle transport, as well as loci known to be associated with retinal diseases. A large fraction of RetMICs and their retina-enriched regulation date back to the last common ancestor of vertebrates, suggesting a critical functional role for this program across vertebrate species. We further identify *SRRM3* as the key regulator of RetMICs inclusion in PRs. Consistently, zebrafish lacking the *srrm3* eMIC domain show progressive OS shortening, PR degeneration and visual impairment concomitant with RetMICs downregulation. Together, these results demonstrate that the conserved program of retina-specific microexons regulated by *Srrm3* is essential for PR functionality and vision, contributing to a novel understanding of retina physiology in health and disease.

## RESULTS

### The human retina has the highest number of tissue-enriched microexons

In order to systematically profile tissue-enriched microexon programs, we used data from *VastDB* (vastdb.crg.eu)^20^ comprising 136 samples from different human cell/tissue types. For each tissue type, we aimed at identifying microexons with biased inclusion, which we define as tissue-enriched (see Methods). We found that the retina has the largest program of tissue-enriched microexons of all analyzed tissues (Fig. 1A). Since many of these microexons are also present in other neural tissues, we implemented a Retina Specificity Score to discriminate events specifically enriched in retina that takes into account the average inclusion level (using the metric Percent Spliced In [PSI]) and its standard deviation for a given exon in retinal, neuronal and non-neuronal samples (see Methods). Using this metric, we found 75 microexons that are specifically enriched in human retinal samples (RetMICs), as well as 116 retina-enriched long exons (i.e. > 27 nucleotides, hereafter RetLONGs) (Table S1). RetMICs were further subdivided into 23 microexons included exclusively in retina (retina-exclusive) and 52 with higher inclusion in retina compared to other neuronal samples (retina-differential) (Fig. 1B).

**Fig. 1 -.**
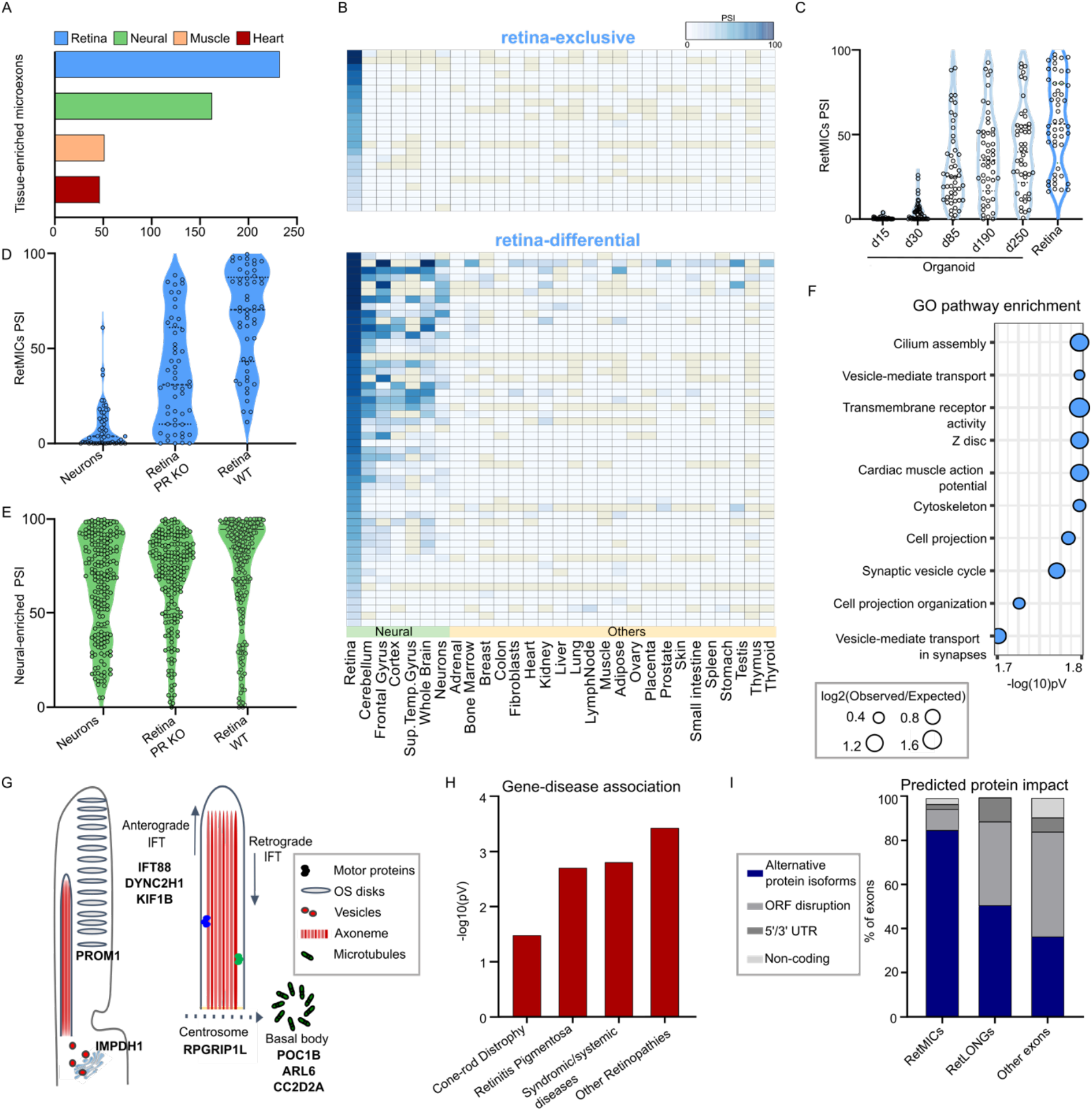
Characterization and identification of the human RetMICs program. **(A)** Number of tissue-enriched microexons by tissue type in humans. Only the four tissues with the highest number of tissue-enriched microexons are depicted. **(B)** Heatmap showing RetMIC inclusion in different human tissues. RetMICs are divided into retina-exclusive (inclusion only in retina samples) and retina-enriched (biased inclusion in retina compared to neural samples). Inclusion levels were obtained from *VastDB^20^*. **(C)** Inclusion levels (PSIs) of RetMICs in cone-rich organoids and whole retina. Developing organoid time points: day 15 (d15), day 30 (d30), day 85 (d85), day 190 (d190) and day 250 (d250). Data plotted are averaged PSIs of three biological replicates; events with NA values due to insufficient read coverage have been omitted. **(D,E)** Violin plots depicting inclusion levels of RetMIC (D) and neural-enriched microexon (neural MICs) (E) in hippocampal neurons, WT and *Aipl1* KO retinas. Events with insufficient read coverage were omitted. **(F)** Top ten enriched GO terms for human RetMIC-containing genes. P-values are FDR adjusted. **(G)** Schematic representation of the OS and localization of selected RetMIC genes; IFT: Intra-Flagellar Transport. **(H)** Enrichment of RetMIC-containing genes among genes associated with different retinal diseases. P-values were calculated with a hypergeometric test. Complete inputs and results are provided in Table S3. **(I)** Predicted protein impact of different exon types as annotated in VastDB.

Although the mammalian retina is primarily composed of PRs, accounting for approximately 60% of all retinal cells^21^, it also contains other neural cell types such as bipolar, ganglion, horizontal and amacrine cells. To determine whether RetMICs are mainly PR specific, we analyzed public RNA-seq data from cone-rich human retinal organoids, developed from human embryonic stem cells and that closely mimic functional cones with mature OS^22^. These data showed that microexon inclusion strongly increases during organoid development and, consequently, PR maturation (Fig. 1C). The inclusion of individual RetMICs across these developmental time-points show strong positive correlations with *Rhodopsin* expression, a marker of PR maturation (median Spearman correlation = 0.83; Fig. S1A-B). Moreover, we analyzed RNA-seq data of retinas from mice knockout for *Aipl1*, which leads to specific PR degeneration^7^. The vast majority (73.5%) of mouse RetMICs (see below) showed a substantial reduction of inclusion (ΔPSI < −15) in *Aipl1* KO mice retina compared to the control (Fig. 1D and Fig. S1C), while neural-enriched microexons did not significantly change their inclusion patterns (Fig. 1E and Fig. S1D). RetMICs also had no or low inclusion in mouse hippocampal neurons, further supporting their PR enrichment (Fig. 1D and Fig. S1C).

### Human RetMICs are enriched in cilia-related genes and associated with retinal diseases

Gene Ontology (GO) analysis revealed that genes containing RetMICs are enriched for functions important for PR homeostasis such as cilium assembly and vesicle-mediated intracellular transport (Fig. 1F and Table S2). We identified RetMIC-containing genes that localize to the cilia basal body and transition zone (*POC1B, ARL6, CC2D2A^23^–^25^*), where they control ciliogenesis and vesicle anchoring to the OS microtubule axoneme. Others regulate OS disks morphogenesis (*PROM1^26^*), PR homeostasis (*IMPDH1^27^*), are involved in anterograde and retrograde transport (*IFT88, DYNC2H1, KIF1B^28,29^*) or are part of the centrosomal complex (*RPGRIP1L^30^*) (Fig. 1G). This contrasts with the GO analysis for genes harboring neural-enriched microexons, which were also enriched in functions related to vesicle-mediated transport and neural development, but not to cilia biogenesis (Table S2). Remarkably, RetMIC-containing genes were significantly enriched among loci associated with multiple retinal diseases (Fig. 1H, Table S3), including Retinitis Pigmentosa, Cone-Rod Dystrophies and Bardet-Biedl Syndrome, pointing to a potential role for RetMICs in vision. In line with this hypothesis, at least two individual RetMICs have been previously linked to visual impairment (*arl6*^31^ and *DYNC2H1*^32^).

### RetMICs generate alternative protein isoforms with remodeled structures

Similar to neural microexons^10^, we found that RetMICs were less likely to disrupt open reading frames than RetLONGs and other cassette exons (Fig. 1I). Moreover, a larger fraction of RetMICs overlapped annotated PFAM or PROSITE protein domains^33,34^ (46% of RetMICs vs 35% of RetLONGs; Table S4). Together, these observations suggest that RetMICs may serve to modulate the activity of protein domains and/or protein-protein interactions, rather than affecting gene expression through nonsense-mediated decay or by causing gross alterations to protein folding. To perform a more comprehensive analysis of the structural impact of RetMIC inclusion, we compiled and examined all available PDB structures of human RetMIC-containing genes as well as high-confidence structural models from ModBase^35^ and Interactome3D^36^ (Fig. S2 and Table S5). Out of 32 protein structures containing the microexon insertion site, the vast majority (96%) of those sites occurred in solvent accessible regions, namely regions with Relative Solvent Accessibility ≥ 20%^37^. Most insertion sites were mapped to unstructured loops (72%) rather than within structured helices (20%), sheets (4%), or turns (4%), further suggesting that RetMICs generally serve to modify protein surfaces instead of impacting overall protein folding.

Through our structural analysis we identified interesting examples of RetMICs in ubiquitous vesicular trafficking-related proteins that are likely to affect their functionality. For instance, a highly retina-specific and evolutionarily conserved 3-nt microexon (VastID: HsaEX0015816) is present in the clathrin heavy chain (*CLTC*) gene, a key protein in charge of packaging cargo into clathrin-coated vesicles at the plasma membrane (Fig. S3A,B). This RetMIC occurs within the N-terminal β-propeller domain, which mediates interactions with multiple clathrin-box motif-containing adapter proteins that compete for binding to the same surface^38^. Upon inclusion, the microexon introduces a single negatively charged aspartic acid residue into the otherwise hydrophobic clathrin-box-binding groove of the β-propeller, thereby modifying the electrostatic landscape of this interaction surface and likely affecting its interaction with adapters (Fig. S3C-E). Another interesting example is the motor protein myosin-VI (*MYO6*), which co-localizes with clathrin-coated vesicles and plays important roles in ciliary vesicular transport^39^. *MYO6* harbors a conserved 9-nt retina-enriched microexon (VastID: HsaEX0041235) in the catalytic motor domain, through which myosin binds to and catalyzes movement along the actin filament through its ATPase activity (Fig S4A,B). The microexon is located within a loop termed “insert-1” that is not shared with other myosins and has been shown to modulate nucleotide binding (Fig. S4C,D)^40^. This insert is thought to explain some of the unique kinetic properties of myosin-VI relative to other myosins, which may be further fine-tuned in PRs by the inclusion of the RetMIC, especially given its highly acidic sequence (Glu-Asp-Glu) that might affect the binding of negatively charged ATP. Another recent study identified a retina-enriched microexon (VastID: HsaEX0021304) within a different motor protein, dynein cytoplasmic 2 heavy chain 1 (*DYNC2H1*), which, from its position within an ATP-binding domain, also appears likely to affect conformational dynamics during microtubule binding and ATP hydrolysis^41^.

### RetMICs are evolutionarily conserved and enriched in cilia-related genes in vertebrates

To investigate the evolutionary conservation of RetMICs across vertebrate species, we used publicly available RNA-seq samples for mouse, chicken and zebrafish^20^ as well as zebrafish adult retina samples that we generated for this study. Similar to humans, retina was the tissue with the largest tissue-enriched microexon program in all species (Fig. 2A). Applying the Retina Specificity Score to define RetMICs in mouse, chicken and zebrafish, we identified 63 mouse, 75 chicken and 72 zebrafish RetMICs (Table S1). To evaluate RetMIC conservation from both the genomic and regulatory perspective, we first derived exon orthologies among all selected vertebrate species using *ExOrthist^42^* (see Methods). RetMICs showed significantly higher levels of genomic conservation compared to RetLONGs (Fig. 2B), with 42%, 62% and 84% of human RetMICs conserved in zebrafish, chicken and mouse, respectively. To evaluate regulatory conservation, we next investigated if genomically conserved RetMICs had retina-enriched inclusion. Preferential inclusion in the retina compared to the other tissues was observed for the majority of the human RetMIC orthologs (Fig. 2C), with few exceptions represented by exons characterized by more broad, neural-enriched inclusion (Fig. 2C and S5). Overall, these results suggest an ancestral role of the RetMIC program in the vertebrate retina. To assess whether RetMICs impact similar biological processes across vertebrates, we then performed comparative GO enrichment analyses. To avoid biases coming from the different genome annotations, we transferred human GO annotations to the mouse, chicken and zebrafish gene orthologs (see Methods). As for humans, cilium organization and vesicle transport appear as enriched categories across the studied species (Fig. 2D), in contrast to RetLONGs (Table S2 and Fig. S5B). In summary, these results show that a program of RetMICs modulating PR ciliogenesis and vesicle transport was likely already present in the last common ancestor of jawed vertebrates.

**Fig. 2 -.**
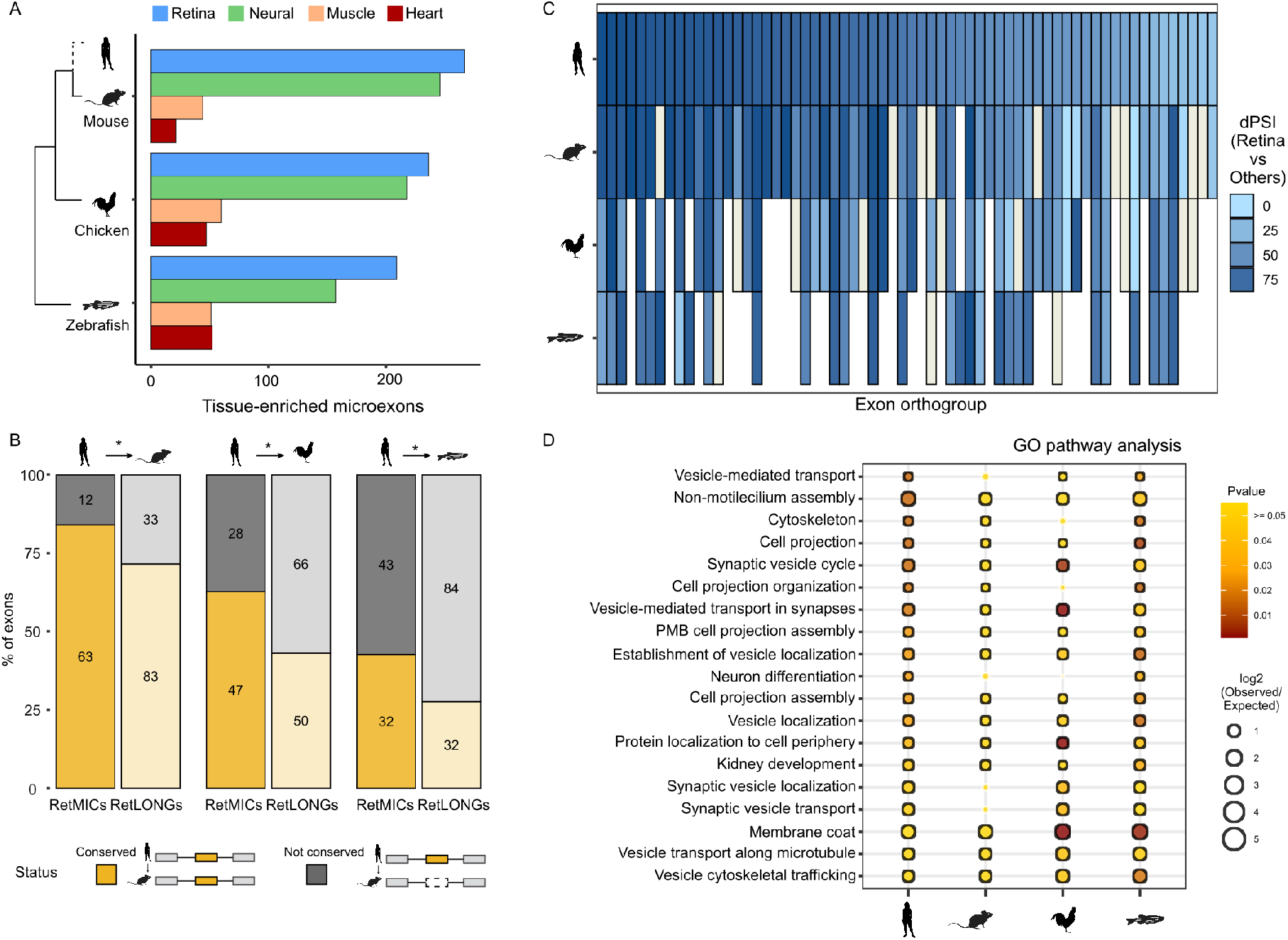
Evolutionary study of RetMICs. **(A)** Number of tissue-enriched microexons (as in Fig. 1A) in mouse, chicken and zebrafish. Only the four tissues with the highest number of tissue-enriched microexons are represented. **(B)** Genomic conservation of human RetMICs (full color, left bar) and human RetLONGs (transparent color, right bar) in mouse, chicken and zebrafish. Human RetMICs/RetLONGs are considered genomically conserved in another species when they belong to an exon orthogroup including at least one exon from that species. Asterisks indicate significant differences between RetMICs and RetLONGs conservation (*P* < 0.05, hypergeometric test). **(C)** Heatmap showing the bias in retina inclusion for genomically conserved human RetMICs (top row) and their respective orthologs in mouse, chicken and zebrafish. The color represents the ΔPSI between the average of the retina samples and of all the other tissues, with darker blue reflecting greater retina inclusion bias. In case of multiple orthologs only the one with the highest ΔPSI (retina-others) was plotted. Blanks and ivory rectangles indicate missing orthologs and missing ΔPSI values due to lack of read coverage, respectively. **(D)** Dot plot representing functional enrichment of RetMIC-containing genes across species. The functional enrichment of genes containing RetMICs was separately tested for each of the species, and significant categories in at least two species (FDR-adjusted *P* ≤ 0.05) were plotted. The color reflects the adjusted p-value, with yellow color depicting *P* ≥ 0.05. The size of the dots is proportional to the log2 of the observed vs. expected ratio (O/E), and black borders around the categories highlight log2 O/E ≥ 1. PMD: Plasma Membrane Bounded.

### *SRRM3* and *SRRM4* regulate RetMIC inclusion *in vitro*

We next looked into the regulation of RetMICs. First, we focused on *MSI1*, a splicing factor that is highly expressed in PRs and can promote the inclusion of retina-enriched exons through direct binding to UAG motifs in their downstream introns^7^. However, stable ectopic expression of *MSI1* for 24 h in HEK293 cells (Fig. S6A) revealed that RetMICs were largely not responsive to this regulator (Fig. 3A,B), in contrast to some RetLONGs and known *MSI1* targets (Fig. 3B). Next, we tested the effect of *SRRM3* and *SRRM4* ectopic expression (Fig. S6A). Similar to neural-enriched exons^10,14^, expression of either gene was sufficient to promote the inclusion of many RetMICs (Fig. 3A,B). Similar results were obtained using previously published data from ectopic expression of *MS1* and *SRRM3/4* in different neural and non-neural cell lines^8,43^. Altogether, 71% of human RetMICs responded substantially (ΔPSI ≥ 15) to *SRRM3/4* expression in at least one experiment, in contrast to 9% for *MSI1* (Table S6). Moreover, *SRRM3/4* and *MSI1* showed the opposite exon length preference: whereas *SRRM3/4* enhanced mostly short retina-specific exons, *MSI1* regulated a larger fraction of long exons than of short exons (Fig. 3C). Analysis of the surrounding intronic sequences revealed a significant enrichment of *SRRM3/4*-binding UGC motifs upstream of RetMICs and, to a lesser extent, of short (28-50 nt) RetLONGs, as compared to longer RetLONGs or a set of random control exons (Fig. 3D). In contrast, both RetMICs and RetLONGs were enriched for *MSI1*-binding UAG motifs in the downstream intron compared to control exons (Fig. 3D). Taken together, these results suggest that *SRRM3/4* activity is sufficient to promote the inclusion of the majority of RetMICs, although *MSI1* may be able to further modulate their inclusion levels.

**Fig. 3 -.**
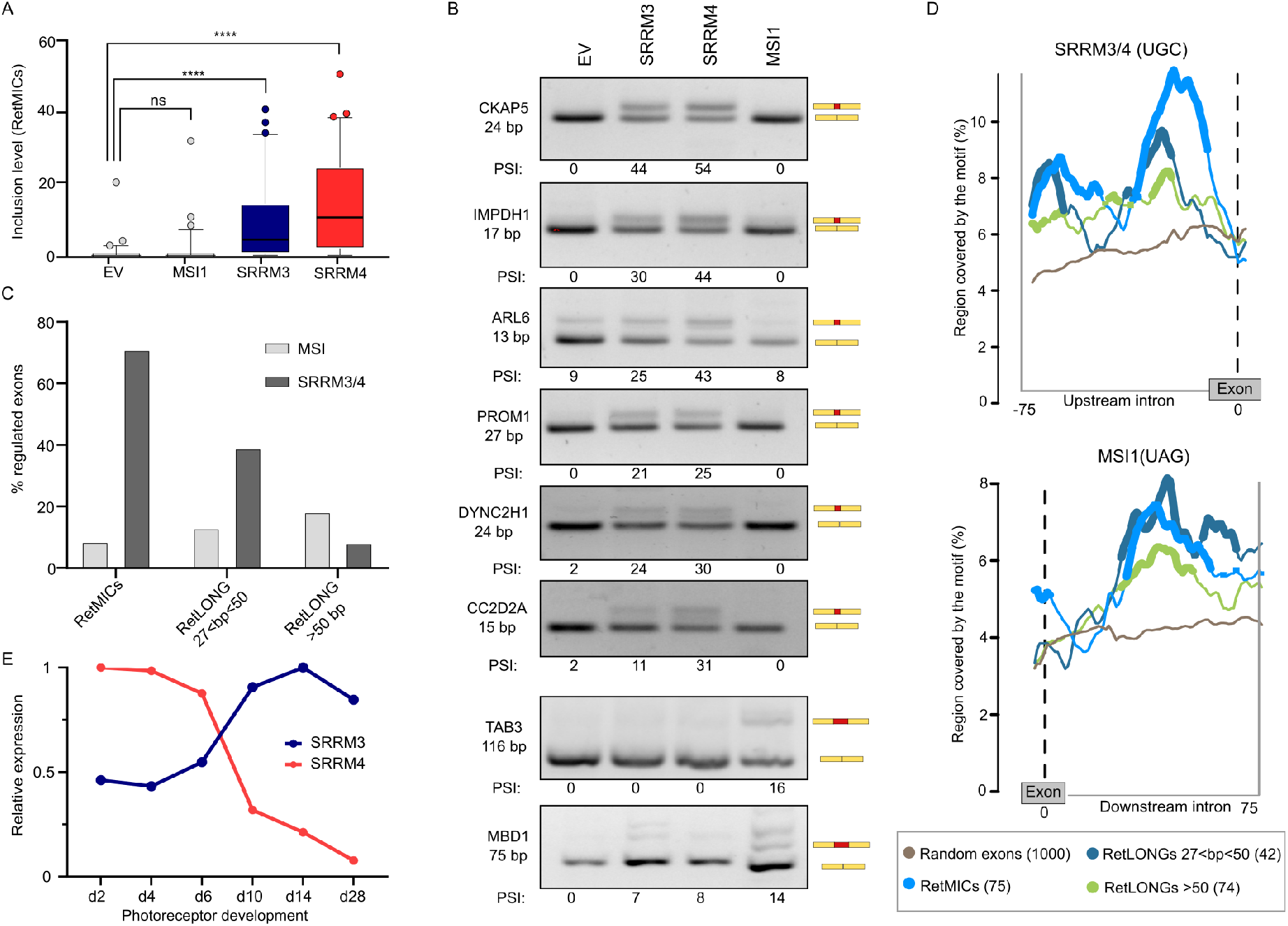
Regulation of RetMICs by *MSI1, SRRM3* and *SRRM4*. **(A)** Inclusion level (PSI) of RetMICs in HEK293 cells ectopically expressing *SRRM3, SRRM4* and *MSI1*. EV: Empty Vector. PSIs are quantified using *vast-tools* on RNA-seq data from cells 24h post-induction with 1 μg/mL doxycycline; **** p<0.0001 Wilcoxon test; NS: not significant. **(B)** RT-PCR assays showing the inclusion of RetMICs (*CKAP5, ARL6, IMPDH1, PROM1, DYNC2H1* and *CC2D2A*), a RetLONG (*MBD1*) and a known MSI1-dependent exon (*TAB3*) in HEK293 cell lines upon ectopic expression of *SRRM3, SRRM4* and *MSI1*. PSI levels quantified using *ImageJ* are shown below each gel. **(C)** Percent of retina-enriched exons by length group that showed substantial upregulation (ΔPSI>15) upon *SRRM3/4* (dark grey) or *MSI1* (grey) expression in at least one experiment. **(D)** RNA maps of SRRM3/4 and MSI1 associated binding motifs in the regions surrounding retina-enriched exons by length group and 1000 random exons. For simplicity, only the relevant upstream (SRRM3/4) or downstream (MSI1) introns are shown. Regions with a significant difference in motif coverage in the tested exon group with respect to random exons (FDR<0.05) are marked by thicker lines. Sliding window is 27 bp. **(E)** *Srrm3* and *Srrm4* gene expression levels (cRPKMs) across mouse developing rods (data from VastDB). Expression levels are relative to the stage with the highest value.

### *srrm3* is necessary for RetMIC inclusion in zebrafish

Unlike in most other types of neurons, *Srrm4* has been shown to be lowly expressed in adult PRs^7^. Indeed, analysis of *Srrm4* expression levels in mouse developing rods revealed a sharp downregulation of its levels over time (Fig. 3E). Interestingly, *Srrm3* displays the opposite pattern, with increasing expression levels during PR maturation (Fig. 3E). This switch in expression from *Srrm4* to *Srrm3* in mature PRs suggests that, while both *Srrm4* and *Srrm3* can induce RetMICs inclusion *in vitro*, *Srrm3* may be primarily responsible for RetMIC inclusion in mature PRs *in vivo*. To evaluate this possibility and investigate the physiological roles of these regulators, we used the CRISPR/Cas9 system to generate zebrafish mutant lines for *srrm3* and *srrm4* (Methods). For each gene we targeted the eMIC domain (Fig. S7A,B), which is necessary and sufficient for microexon regulation^14^. While fish homozygous for the *srrm4* mutation (*srrm4* MUT) did not display any evident phenotype, including changes in size or survival rate, *srrm3* MUT and double homozygous mutant (DMUT) larvae died between 10 and 13 days post fertilization (dpf) (Fig. 4A). We hypothesized that this early mortality was linked to visual impairment, as reported for other zebrafish models of blindness in which the mutant fish are unable to forage for food and die of starvation after exhaustion of the yolk sac^44^. If this were the case, we would expect that, in dark conditions, all genotypes resulting from a heterozygous (HET) cross would be equally affected, resulting in genotype ratios consistent with the Mendelian expectation. In line with this, in contrast to the specific depletion of homozygous mutants observed in control light conditions, the fish that survived at 13 dpf in the dark showed no homozygous depletion (26% WT, 39% HET, and 35% MUT) (Fig. 4B). These results thus indicate that, whereas all genotypes are equally likely to die in darkness, only MUT fish are more likely to do so in light conditions, pointing to a visual impairment upon *srrm3* depletion.

**Fig. 4 -.**
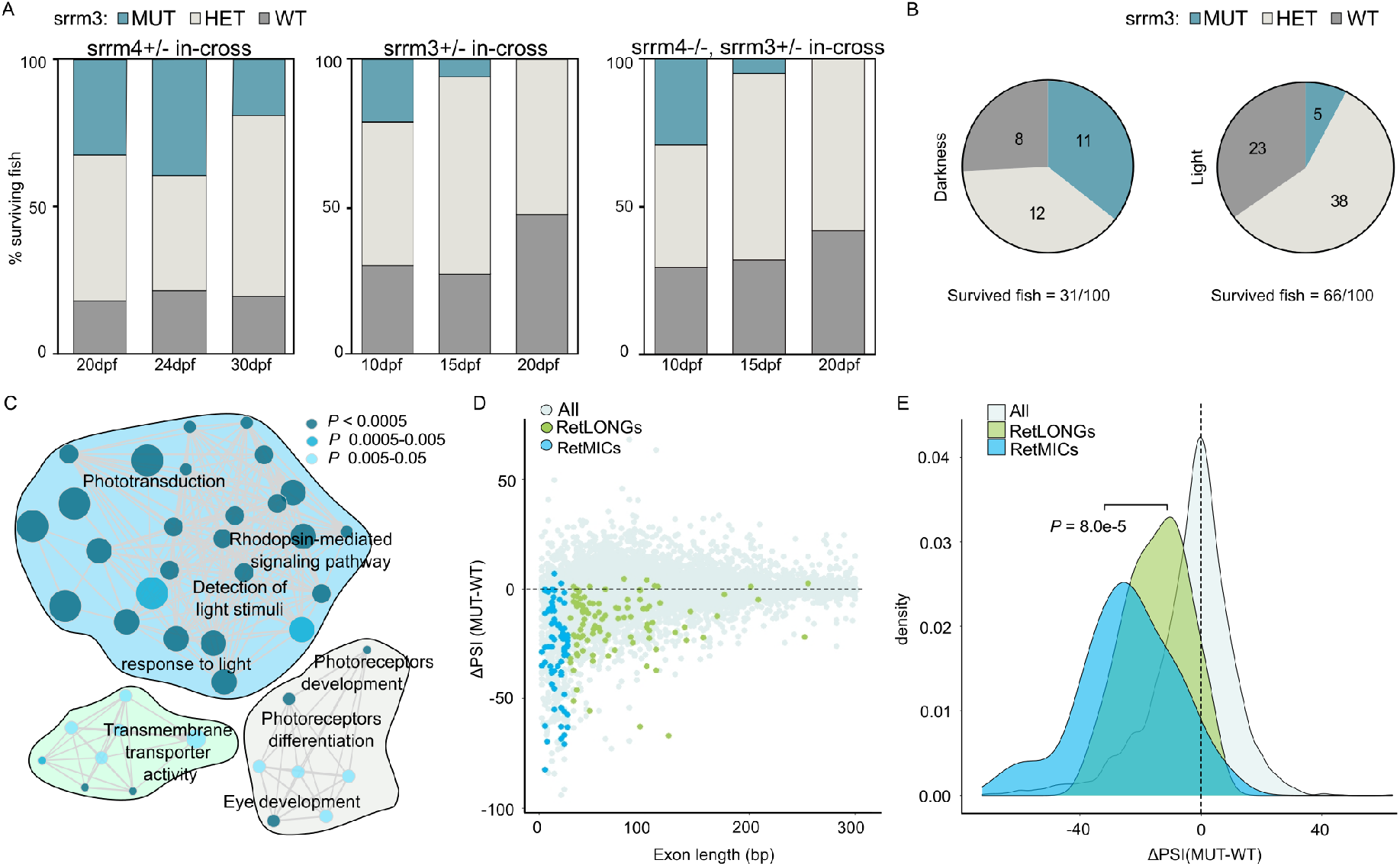
*srrm3* depletion in zebrafish causes early lethality and RetMIC downregulation. **(A)** Genotype distribution for surviving larvae at different time points from in-crosses of *srrm4+/-, srrm3+/-* or *srrm4-/-;srrm3+/-* (double mutants) fish; dpf: days post fertilization. **(B)** Genotype distribution for surviving larvae at 13 dpf from an *srrm3+/-* in-cross in dark and control light conditions. **(C)** Enriched Biological Process GO terms for genes downregulated in *srrm3* homozygous mutants (MUT) eyes (log2FC(MUT/WT) ≤ −1.5). GO terms are grouped by ClueGO into three networks according to their GO Groups. GO Groups are highlighted using three different arbitrary colors, as listed in Table S7. P-values are corrected with Bonferroni step down. **(D)** Change in inclusion levels (ΔPSI (MUT-WT)) quantified using *vast-tools* for all exons shorter than 300 bp with sufficient read coverage in WT and *srrm3* MUT eyes. Blue/green dots correspond to RetMICs and RetLONGs, respectively. **(E)** Density plots for ΔPSI distributions of RetMICs, RetLONGs and other alternative exons (10<PSI<90 in WT and MUT) (*P* = 8.0e-5; Wilcoxon Rank-Sum test).

To investigate the effect of the *srrm3* depletion in the retina at the molecular level, we next examined gene expression and splicing changes by enucleating the eyes of WT and *srrm3* MUT fish at 5 dpf and performing RNA-seq. Several genes crucial for PR functionality showed reduced expression in *srrm3* MUT eyes (e.g. *rhodopsin*: log2FC(MUT/WT) = −3.72) (Table S7), and GO analysis of downregulated genes further revealed a strong enrichment for visual function and phototransduction (Fig. 4C and Table S7). Quantification of alternative splicing events using *vast-tools* revealed multiple mis-regulated exons, 53% (225/409) of which corresponded to microexons. Moreover, although both RetMICs and RetLONGs showed global downregulation in mutant eyes (Fig. 4D and Table S1), RetMICs exhibited significantly larger decreases in inclusion levels than RetLONGs (*P* = 8.0e-5; Wilcoxon Rank-Sum test) (Fig. 4E).

### *srrm3* is necessary for OS maintenance and visual function in zebrafish

To follow up on these observations pointing at major visual defects, we then investigated the retinal morphology of the mutants. We performed immunostaining of frozen retinal sections of 5 and 10 dpf larvae using: (i) ZPR-3, a PR marker commonly used to stain Rhodopsin in the OS^45^, and (ii) ZPR-1 (Arrestin3), a PR-specific antigen expressed in red and green double cones^45^. ZPR-3 staining at 5 dpf revealed that OSs appear severely shortened and disorganized in *srrm3* MUT larvae, with different spots corresponding to Rhodopsin mis-localization throughout the outer nuclear layer (ONL) (Fig. 5A,B). Moreover, we observed a significant decrease of the ONL thickness in mutant retinae (Fig. 5C and Table S8), a common signature of retinal diseases preceding PR degeneration. Indeed, at 10 dpf only a few spots of Rhodopsin were detected in the mutants, while the ONL disappeared completely (Fig. 5D-F). ZPR-1 staining confirmed that cones also displayed progressive degeneration and mis-localization to the cell bodies at 5 and 10 dpf (Fig. S7C). The DMUT retinal phenotype at 5dpf was similar to that of *srrm3* MUT but stronger (Fig. S7D). In contrast, histological examination of single *srrm4* MUT retinae revealed no morphological changes compared to WT fish (Fig. S7C).

**Fig. 5 -.**
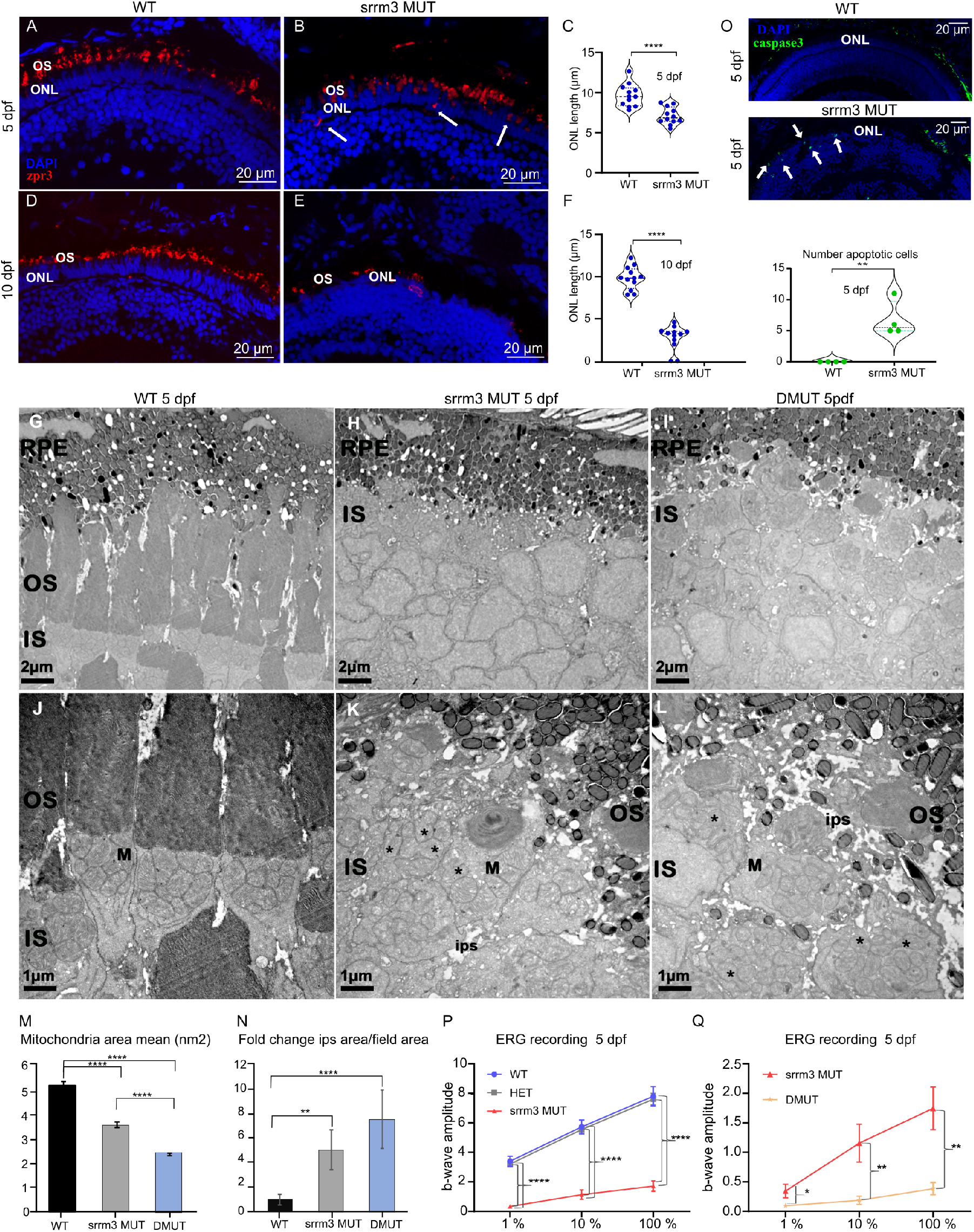
Photoreceptor degeneration and visual impairment in *srrm3* mutants. **(A-F)** ZPR-3 staining in *srrm3* WT (A and D) and *srrm3* homozygous mutants (MUT) (B and E) at 5 and 10 dpf. Arrows show Rhodopsin mis-localization. N = 5 for 5 dpf and N = 3 for 10 dpf fish. Quantifications of the thickness of the ONL are provided for 5 dpf (C) and 10 dpf (F) as measured in three areas of the retina, one section per animal. N = 4, p-values correspond to unpaired t-test. (G-N) Electron microscopy images show absence of OSs or a dramatic OS length decrease in eyes of both *srrm3* MUT (H-K) and double homozygous mutants (DMUT) (I-L) compared to WT ones (G-J) at 5 dpf. Further magnification (J-L) revealed smaller mitochondria (asterisks indicate mitochondria in fission process) and enlarged interphotoreceptor space (ips), quantified as mean of mitochondria area per field and ips area normalized on field area, represented as fold change in M and N, respectively. One-way ANOVA test with Tukey post-hoc analysis was applied. Scale bars are 2 μm in F-H and 1 μm in I-M. *N* ≥ 4 eyes/genotype. N ≥ 13 of fields/genotype was analyzed. (P,Q) ERG responses from WT, HET, and *srrm3* MUT (P) or *srrm3* MUT and DMUT (Q) at 5 dpf upon different light stimuli (1%, 10% and 100%). All recordings were done in two biological replicates. N = 20 for WT, N = 34 for HET, N = 17 for *srrm3* MUT (P), N = 22 for *srrm3* MUT (Q), and N = 15 for DMUT. P-values from one-way ANOVA. (O) Caspase-3 staining and associated quantifications in 5 dpf retina sections. N = 4. Significance code: ****, *P* = 0; ***, 0 < *P* < 0.001; **, 0.001 ≤ *P* < 0.01; *, 0.01 ≤ *P* < 0.05.

We next used electron microscopy to analyze OS ultrastructure. This analysis confirmed that OSs were dramatically shortened and disorganized at 5 dpf in both *srrm3* MUT and DMUT eyes (Fig. 5G-L), indicating that the ZPR-1 and ZPR-3 signals observed by immunofluorescence largely corresponded to mis-localized proteins. These results also revealed smaller mitochondria in both mutants (Fig. 5J-L), with a decrease in the total mitochondria area per field with respect to WT retinae (Fig. 5M and Table S9). In particular, the mitochondria appeared more fragmented in mutant eyes, indicating an increase in the fission process (asterisks in Fig. 5K,L), which represents an early event prior to PR cell death^46,47^. This phenotype was more severe in DMUTs, showing increased numbers of mitochondria, which are smaller and even more fragmented, probably due to a higher rate of fission (Fig. S7E and Table S9). Mutant retinae also exhibited an enlargement of the interphotoreceptor space, particularly in DMUT eyes (ips, Fig. 5N, Table S9). This interphotoreceptor space enlargement has been reported to occur due to the accumulation of extracellular vesicles^48^ and is commonly observed in animal models of PR degeneration caused by alterations in ciliary trafficking^49,50^, suggesting the presence of compromised PR cells. To monitor PR cell death, we stained retinal sections from WT and *srrm3* MUT larvae at 5 dpf with an anti-caspase 3 (active caspase 3) antibody. The number of caspase 3-positive cells was significantly increased in MUT animals compared to WT siblings, with the signal being specifically localized to the PR layer (Fig. 5O and Table S8).

Altogether, these results point to strong defects in retinal function. To directly monitor visual performance, we did electroretinography (ERG) recordings for 5 dpf larvae (Table S10). ERG is a non-invasive test measuring field potential change of the whole retina induced by light^51^, resulting in characteristic waveforms defining the neurons contributing to the response. In the zebrafish retina, photoresponses are dominated by cones^52^ and they are represented by a b-wave of a certain amplitude. We observed that *srrm3* MUT fish have a striking decrease in b-wave amplitude upon light stimulation with respect to HET and WT siblings, indicating a strong visual impairment (Fig. 5P). Moreover, DMUT fish performed even worse than *srrm3* MUT fish (Fig. 5Q), supporting a minor but significant overlapping role for *srrm4*. Nevertheless, single *srrm4* MUT fish did not display any significant visual impairment compared to WT siblings (Fig. S7F). Taken together, these results confirm that *srrm3* is essential for proper microexon inclusion in the zebrafish retina as well as for PR maintenance and responsiveness to light stimuli and vision.

## DISCUSSION

Alternative splicing is an essential mechanism for generating molecular and functional diversity across cell and tissue types. In the context of the nervous system, which shows a particularly high degree of tissue-enriched alternative splicing, PRs differ in their transcriptomic profiles from other neuronal subtypes^53^, likely as a reflection of their unique cellular morphology and function. Our results reveal a novel program of retina-enriched microexons that further contribute to this molecular specialization. These microexons are included in PRs in addition to neural microexons, which are shared with other neuronal types, and both microexon programs have distinct and common regulatory and functional properties.

We demonstrate that the ectopic expression of either *Srrm3* or *Srrm4* is sufficient to drive the inclusion of most RetMICs in non-PR cells, as it was shown for neural microexons^10,19^. However, we show that only *Srrm3* is highly expressed in mature PRs and that the inclusion of most RetMICs depends mainly on this paralog *in vivo*. Nevertheless, we also found evidence for a minor but significant redundant role for *srrm4* in the retina, since double mutant fish displayed stronger visual impairment and histological defects than single *srrm3* mutants. Given the substantial levels of both *Srrm3* and *Srrm4* expression in other neuronal types^19,54^, these results thus raise the question of why RetMICs, particularly the retina-exclusive ones, have low or no inclusion in non-PR neurons. At least two non-mutually exclusive hypotheses may explain this pattern. First, although we found that the PR-specific splicing factor MSI1 on its own is not sufficient to drive the inclusion of most RetMICs, it is possible that MSI1 is necessary for RetMIC splicing in PRs *in vivo*, acting synergistically with SRRM3. In line with this hypothesis, we found a significant enrichment for MSI1 binding motifs in the downstream introns of both RetMICs and longer retina-enriched exons, a binding location that is expected to promote exon inclusion^7^. Second, PRs are known to have very low expression of some major neural splicing factors, including members of the Nova, Rbfox and Elavl families (Fig. S8A)^53,55^. Therefore, it is possible that these splicing factors act as negative regulators of RetMICs in non-PR neurons, allowing for their inclusion only in PRs. Consistently, we found a significant enrichment for known binding motifs for Nova, Rbfox and Elavl in the upstream introns of RetMICs (Fig. S8B), where their binding is expected to cause exon downregulation^56^. This is in stark contrast with neural microexons, which showed enrichment for inclusion-enhancing Nova and Rbfox binding motifs in the downstream introns (Fig. S8B), as previously reported ^11,57^. Further research should evaluate these hypotheses to provide a more complete mechanistic understanding of the unique inclusion profile of RetMICs promoted by *Srrm3*.

At the functional level, we found that, similar to neural microexons^10^, RetMICs can remodel protein structures of genes enriched for vesicle-mediated transport and related functions. However, we also revealed an enrichment for genes involved in cilium assembly, which was not observed for neural microexons. Altogether, these functions suggest that RetMICs enable a unique proteome specialization that is necessary for proper development and functioning of the OS, the highly modified cilium of PRs. Given the unusually high demand for vesicle formation, transport, and recycling in the OS^4^, the specific modifications introduced by RetMICs in otherwise ubiquitous trafficking and ciliary machinery may help meet this high demand by promoting interactions with PR-specific substrates and/or facilitating unique catalytic properties not required (or even detrimental) in other cell types. In line with this idea, depletion of *srrm3* caused severe malformation of the OS and vesicle accumulation, leading to PR degeneration and impaired visual function. Importantly, in homozygous *srrm3* mutant larvae, OSs are generated but not maintained. This is similar to what has been described for various zebrafish mutants of terminal effector genes involved in intraflagellar transport^58^, where mis-localization of visual pigments is associated with OS disappearance and PR degeneration. Furthermore, mutation of other genes necessary at different stages of ciliogenesis or for vesicle trafficking^50,26,59^, result in PR phenotypic alterations characterized by vesicle accumulation and dysmorphic OSs. Remarkably, the nearly complete loss of the OSs that we report for *srrm3*, places this microexon regulator among the genes with the strongest mutant PR phenotypes reported so far in zebrafish, with features matching the ones observed in human Retinitis Pigmentosa^60^.

It should be noted, however, that depletion of *srrm3* also affected the inclusion of many neural-enriched microexons and other short exons in zebrafish eyes, whose mis-regulation certainly contribute to the observed phenotypes. Nevertheless, three additional observations support that RetMICs are likely to be directly behind at least some of the *srrm3* mutant phenotypes. First, the inclusion of RetMICs was particularly mis-regulated in the eyes of *srrm3* mutants. Second, as mentioned above, mis-regulation or mutation of the RetMICs in *arl6* ^31^ and *DYNC2H1* ^32^, respectively, have been shown to directly impact visual function in line with the *srrm3* mutant phenotypes. Third, zebrafish lacking individual RetMIC-containing genes, such as *cc2d2a* ^50^, *ift88* ^58^ or *prom1b* ^26^, have been shown to display disorganization of the vesicle fusion machinery and strong PR degeneration. Therefore, it is plausible that the multiple PR-related phenotypes caused by *srrm3* depletion are at least in part caused by mis-regulation of RetMICs. Remarkably, this would place RetMICs as novel candidates to underlie retinopathies without a known genetic cause, opening new avenues for the understanding of the highly complex molecular genetics of retinal diseases.

## METHODS

### Definition of tissue-enriched microexons

We downloaded information for all alternative splicing events of all types from *VastDB^20^* (https://vastdb.crg.eu/wiki/Downloads) including their inclusion levels (using the Percent Spliced In [PSI] metric as provided by *vast-tools*) across all *VastDB* samples and their genomic features (e.g. length, sequence, genomic coordinates, predicted protein impact) for all the species included in the manuscript (human: hg38, mouse: mm10, chicken: galGal4, zebrafish: danRer10). We considered as microexons all cassette exons with length ≤ 27 nucleotides. We then used the *Get_Tissue_Specific_AS.pl^61^* script (https://github.com/vastdb-pastdb/pastdb) to derive sets of tissue-enriched microexons from PSI tables. We defined as tissue-enriched in a tissue T all microexons presenting: (i) ΔPSI (delta PSI) between tissue T average (i.e. the PSI average among all samples from tissue T) and each of the other tissues’ averages ≥ 15, (ii) ΔPSI between tissue T average and the average of all the other tissues ≥ 25, (iii) coverage in at least N tissues (with N=10 for human and mouse; N=8 for chicken and zebrafish), and (iv) at least n samples per tissue (with n=2 for human and mouse; n=1 for chicken and zebrafish). In order to capture microexons with biased inclusion patterns in one or more tissues, for some tested tissues (e.g. Neural) we excluded tissue groups known to be characterized by partially overlapping microexon programs (e.g. Retina, Muscle, Heart) from the pool of compared tissues. We built a config file for each species, highlighting the correspondence between sample, relative tissue group and tissue groups excluded from the comparison, which we provided as input to *Get_Tissue_Specific_AS.pl* script. The config files used in the manuscript are listed in Table S11.

### Definition of RetMICs and RetLONGs

In order to identify microexons with retina-specific inclusion biases (i.e. enriched in no other tissue), we ran *Get_Tissue_Specific_AS.pl* as described in the previous section but without excluding any tissue groups from the pool of compared tissues (config files in Table S11) and printing out a table containing all the ΔPSI Retina-Others or Neural-Others (*--test_tis Retina Neural* option) for all the events with minimum coverage and number of replicates (see (iii) and (iv) in the previous section). We then devised a Retina Specificity Score (RSS) to help the identification of retina-specific exons (RetMICs and RetLONGs), and is defined as follows:

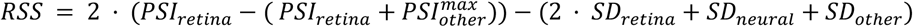

where:

PSI_[Retina|Neural]: exon average PSI among all [retina|neuronal] samples.
SD_[Retina|Neural|Other_tis]: standard deviation of exon PSI distributions across all [retina|neuronal|non-retina] samples.
Max_Other_tis: maximum average PSI among all non-retina tissues (in most cases, Max_Other_tis = PSI_Neural).

We identified RetMICs and RetLONGs for all species (human, mouse, chicken and zebrafish) requiring either (i) ΔPSI between retina and the tissue with the highest PSI ≥ 40, or (ii) average ΔPSI between Retina and all other tissues ≥ 10 and the RSS > 0. We then further divided RetMICs between microexons either exclusively included in retina (“retina-exclusive”; Max_Other_tis ≤ 4 || (4 < Max_Other_tis <15 && PSI_Retina ≥ 90)) or presenting a substantially higher degree of retinal versus neural inclusion (“retina-differential”; all the other RetMICs).

### Exon orthology calls

We ran *Broccoli* (v1.0)^62^ with default parameters to infer gene orthogroups between human (hg38), mouse (mm10), chicken (galGal6) and zebrafish (danRer10), and we selected all gene orthogroups containing 20 genes or less (16,166 out of 16,313). We then ran *ExOrthist main* (v0.1.0)^42^ to derive genome-wide, multi-species exon orthogroups among all species. We considered the following species pairwise evolutionary distance ranges [short: human-mouse (hg38-mm10), human-chicken (hg38-galGal6), mouse-chicken (mm10-galGal6); medium: human-zebrafish (hg38-danRer10), mouse-zebrafish (mm10-danRer10), chicken-zebrafish (galGal6-danRer10)] and set the evolutionary conservation cut-offs to the default values. In order to increase the sensitivity of the exon orthology calls, we also considered non-annotated exons identified by *vast-tools^20^* (*--extraexons* option) and we provided pre-computed *liftOver* hits among each pair of species (computed with the *ExOrthist* companion script *get_liftovers.pl)* to be directly integrated in the exon orthogroups (*--bonafide_pairs* option). We identified a total of 206,644 exon orthogroups, recovering from a minimum of 77.89% of exons (chicken) to a maximum of 88.34% (mouse). Human RetMICs/RetLONGs exons were considered as genomically conserved in one of the other species when belonging to an exon orthogroup including at least one exon from that species (see Fig. 2B).

### GO enrichment analysis and hypergeometric tests

All GO enrichment analyses were carried out with the *gprofiler2* R package^63^, to which we provided custom annotations and backgrounds. We downloaded the human annotation (geneID-GO correspondence) from Ensembl (v102), and we combined it with the clueGO v2.5.5^64^ human annotation for biological processes (level 5) in order to include possibly unique annotations from the two sets. For the evolutionary comparison and to avoid biases due to different annotation qualities between species, we derived annotations for mouse, chicken and zebrafish starting from the assumption that orthologous genes likely share functional properties. We considered the gene orthogroups inferred between human, mouse, chicken and zebrafish (see “Exon orthology calls”) and assigned the functional annotation of each human gene to all the genes from the other species belonging to the same orthogroup. Then, for each species, we filtered for the GO categories with a number of genes included between 5 and 3,500. Background sets for each species consisted of genes containing events of any kind with the expression level requirements used to define tissue-enriched exons (see above). In particular, they were derived by running the *Get_Tissue_Specific_AS.pl* script with the same parameters used for the tissue-enriched microexon call and requiring sufficient read coverage in Retina, but for all event kinds (*--event_type* option set to EX, IR, Alt3, Alt5, in four different runs). The total number of background genes was 14,471 in human, 11,586 in mouse, 11,994 in chicken and 14,588 in zebrafish. False Discovery Rate (FDR) correction was applied to all performed tests. All the enriched GO categories are listed in Table S2.

For visualization purposes, GO enrichment analysis for genes downregulated (condisering only the genes with log2FC(MUT/WT)< −1,5) upon *srrm3* depletion in zebrafish eyes was performed directly using the software ClueGO^64^, selecting Biological Process as ontology. For this purpose, we converted the list of downregulated genes into human orthologs using DIOPT (DRSC Integrative Ortholog Prediction Tool)^65^ with a score ≥ 2 and “best score” required, as previously described^66^. All the enriched GO categories are listed in Table S7.

All hypergeometric (Fisher) tests mentioned in the manuscript were performed through the *phyper* function in baseR. The total number of genes used to compute the hypergeometric input values were derived from the background employed for the GO enrichment analysis (see previous section). Gene-disease information on inherited retinal disease comes from the database RetNet (https://sph.uth.edu/retnet/). The input values and resulting p-values for all the tests are provided in Table S3.

### Structural analysis of RetMICs

A list of 75 human genes and their relative microexon events was used to retrieve the corresponding SwissProt (manually annotated) protein identifier (ID). The gene AC007246.3 was found to correspond to an antisense RNA and MAGI2-AS3 to a non-coding RNA. Protein Data Bank (PDB) structures were downloaded for those SwissProt IDs having any. For all 75 genes (excluding 7 causing ORF disruption), interaction models were searched within Interactome3D^36^ and ModBase^67^, respectively collecting protein-protein interactions (both experimental and modeled) and protein homology models. Redundant experimental interactions, already found in PDB structures, were excluded. Structures collected in ModBase came with a set of submodels with different quality levels and protein coverage. For each gene, the selected submodel was the one presenting the highest set of scores indicated by the authors^35^, with the condition of containing the microexon event and being based on a template structure with at least 40% of sequence similarity. Flanking exons were translated to their amino acid sequences and searched within their corresponding SwissProt entries. Four exon residues, flanking the microexon event, were used to match and determine the event position within collected structures. The structural characterization is available in Table S5. A graphical overview in Fig. S2 shows the events locations captured in 3D structures.

### Regulatory analysis of RetMICs and RetLONGs

HEK293 lentiviral cell lines used in this study to ectopically express *SRRM3, SRRM4* or *MSI1* were generated using a previously described protocol^43^. The gateway destination vector pCW57.1 containing SRRM4 was generated as described previously^43^. SRRM3 and MSI1 cloned into the donor vector pDONR223 were obtained from the Biomolecular Screening & Protein Technologies Facility at CRG. The resulting entry clones were reacted with pCW57.1 using LR clonase II (11791020, Thermo Fisher Scientific). pCW57.1 lentiviral vector (empty vector (EV)) was used as a negative control. Prior to harvesting RNA, expression of the transgenes was induced using 1 μg/mL of Doxycycline for 24 h. RNA was extracted with the RNeasy Mini kit (QIAGEN). Standard polyA-selected Illumina libraries were generated at the CRG Genomics unit and were sequenced in a Illumina HiSeq2500, producing an average of ~80 million paired-end 125 nt reads. Mapping statistics are provided in Table S13. Detection of microexons and longer exons upon induction was done by PCR using Q5^®^ High-Fidelity DNA Polymerase kit (New England biolabs), according to the manufacturer’s instructions. Primers were obtained directly from VastDB, which provides automatically designed primers annealing to the flanking upstream and downstream constitutive exons. PCR products were run on 3.5% agarose gel in TBE buffer to allow small isoform separation. All primer sequences are shown in Table S12.

To assess the enrichment of sequence motifs associated with *SRRM3/4* (UGC) and *MSI1* (AUG) regulation in the upstream and downstream RetMIC and RetLONG intronic sequences, we generated RNA maps running the *rna_maps* function from *Matt^68^* using sliding windows of 27 nucleotides and 75 nt upstream/downstream as region surrounding the exons. To evaluate the enrichment of *NOVA* (YCAY), *RBFOX* (GCATG) and *ELAVL* (TTTNTTT) binding for RetMICs and neural microexons (Fig. S8) we employed a similar approach but considering 150 nt as the region surrounding the exon. For estimating FDR-corrected p-values in the RNA maps we used a permutation test using 1,000 permutations and a threshold of FDR ≤ 0.05 as implemented in *Matt*.

### Generation of *srrm3* and *srrm4* zebrafish mutant lines

Fish procedures were approved by the Institutional Animal Care and Use Ethic Committee (PRBB-IACUEC). Zebrafish (*Danio rerio*) were grown in the PRBB facility at 28°C 14 h light/10 h dark cycle. Individual and massive crosses were carried out in crossing tanks and the resulting eggs were screened for overall health from embryo collection to 5 dpf. Embryos were collected into Petri dishes with E3 medium with methylene blue and placed into the incubator. Sibling WT and mutant larvae between 5 and 10 dpf were used for most of the procedures and WT adult fish of AB background for retina dissection. To create zebrafish mutant lines we used a CRISPR-Cas9 strategy adapting a previously described protocol^69^. Single or double guide RNAs (gRNAs) were designed using CRISPRscan^70^ to target the eMIC domain of *srrm3* (Tg(HuC:GFP; srrm3_eMIC)) and *srrm4* (Tg(HuC:GFP; srrm4_eMIC)) and were injected at 1-2 cell embryos stage. Selected gRNA sequences are: GGGAATAACTGCGTGAGCGGCGG (*srrm3*) and GCTGTGCTTTCTGCTCTTGCAGG and TGATTCTGCGGGCTTCCAGGTGG (*srrm4*). CRISPR-Cas9 injection mix (with a final volume of 5ul) consisted of Cas9 protein (300ng/μl) - PNABio Cas9 protein (CP01-50), the in-vitro transcribed gRNAs (50ng/μl). Additionally, we included 0.5 ul Phenol Red and sterile water to reach 5ul final volume. We injected 1 nl of CRISPR-Cas9 mix in each embryo. For each gene, we generated two independent founders. The screen for germline mutations of *srrm4* at F0 revealed a founder containing a deletion of 1bp upstream the eMIC domain that produces a frameshift mutation causing a premature STOP codon breaking the eMIC domain (Srrm4_F1). The screen for germline mutations for *srrm3* revealed a founder for *srrm3* eMIC MUT line containing a 5nt deletion that altered the reading frame upstream the eMIC, producing a non-functional domain and altering the C-terminal region of the protein (Srrm3_F1). Additionally, we generated a second founder for *srrm3* carrying a 19 bp deletion (Srrm3_F2) and a second founder for *srrm4* carrying 11 bp deletion, both confirming the gross phenotype (Srrm4_F2). All the experiments performed in this manuscript have been done using the Srrm3_F1 and Srrm4_F1 founders but the second founders show similar phenotypes. Founders were crossed with WT strain and F1 animals were genotyped. Heterozygous animals were incrossed to generate the single mutant lines. To obtain the DMUT line, *srrm3* and *srrm4* heterozygous mutants were crossed and *srrm3+/-,srrm4-/-* fish were maintained as viable and fertile. All experiments using DMUTs in this study were done through in-crosses of *srrm3+/-,srrm4-/-* fish, since *srrm3* and *srrm4* are linked in zebrafish making it impractical to use HET/HET incrosses (1/100 DMUT embryos). Therefore, *srrm4* MUT fish were used as sibling controls for DMUT embryos. For fish genotyping, a piece of the caudal fin was cut and disaggregated using 100 uL of NaOH 50mM for 15 minutes at 96 degrees. To neutralize the reaction, 10uL of Tris-HCl pH=7.4 was added. 2 ul were used as templates for PCR with the *srrm3* or *srrm4* primers designed to amplify the genomic region of interest (listed in Table S12) and GoTaq^®^ Flexi DNA Polymerase kit (Promega).

### Isolation of zebrafish larval eyes and adult retinae

For retina isolation, adult animals were euthanized in 0.08% tricaine and whole eyes were removed. Retina isolation was performed as previously described^71^. Five retinae were pulled for RNA extraction. For larval eye isolation, WT and MUT larvae at 5 dpf were euthanized in 0.08% tricaine and whole eyes were removed. Twenty eyes per genotype were pulled for RNA extraction. For that purpose, we used fresh tissues or frozen tissues stored at −80 degrees in RNAlater (Sigma-Aldrich). Tissues were homogenized in RLT Buffer using a BeatBeater. RNA was then extracted using the RNeasy Mini kit (QIAGEN), according to manufacturer’s instructions. For tissue extraction, beta-mercaptoethanol was added to the RLT buffer, following manufacturer’s instructions. RNA for each sample was then used to generate standard polyA-selected Illumina libraries at the CRG Genomics unit that were sequenced in a Illumina HiSeq2500, producing an average of ~125 million paired-end 125 nt reads. Mapping statistics are provided in Table S13.

### Immunofluorescence and histological retinal analyses

Zebrafish eyes were fixed in 4% paraformaldehyde (PFA) over-night, cryoprotected with 30% sucrose overnight, embedded in OCT and cryosectioned. Twenty-micrometer cryosections were collected on slides. For zebrafish staining, sections were blocked in 1% bovine serum albumin, 0.5% Triton X-100 in PBS for 1 h RT and incubated with primary antibodies (ZPR-1 1:400 and ZPR-3 1:200, Zebrafish International Resource Center (ZIRC)) overnight in blocking solution. Sections were then incubated with the Alexa Fluor secondary antibodies (1:1000; Invitrogen) and counterstained with DAPI (Vector Laboratories).

ONL thickness was manually measured by the *ImageJ* tool in three different areas of the WT and MUT retina at 5 and 10 pdf (1 retina section for each eye). Unpaired t-test was applied to measure significance between groups and conditions. The Caspase-3 assay was performed on retina sections following the protocol described above (anti-caspase 3 (Fisher Scientific, 15889738) 1:500). The staining of positive control, WT and MUT retinae was done in parallel. Apoptotic cells were manually counted on the entire ONL in 1 section for each eye by the ImageJ tool. Unpaired t-test was applied to measure significance between groups. For all the experiments, the number of animals/genotypes is indicated in each figure legend.

### Electron Microscopy

Larvae at 5 dpf have been euthanized in 0.08% tricaine, genotyped and fixed using a mixture of 2.5 % glutaraldehyde and 2 % paraformaldehyde prepared in 0.2 M HEPES buffer for 24 h at 4 °C. After fixation, the samples were washed 3 times with PBS 1X. The samples were postfixed in 1% osmium tetroxide, dehydrated, and embedded in epoxy resin. The samples were cut on ultra-microtome Leica EM UC7 and collected on the single-slot oval grids and analyzed with FEI electron microscope. OS, mitochondrial and interphotoreceptor space area was determined using FEI software. The analysis was carried out on 4 eyes/genotype, for each genotype an N ≥ 13 fields was analyzed. One-way ANOVA test was applied for mitochondria Electron Microscopy analysis. As the statistical test is significant (p-value < 0.05), a post-hoc analysis based on Tukey test was applied to determine pairwise comparisons between condition levels with correction for multiple testing (significance codes: p < 0.0001 ‘****’, < 0.001 ‘***’, < 0.01 ‘**’, < 0.05 ‘*’). To test the data are normally distributed we used Shapiro-Wilk test on the ANOVA residuals (i.e., if the *p*-value > 0.05, then the normality is not violated), while we applied Levene’s test to verify the variance across groups is homogeneous (i.e., if the *p*-value > 0.05, then we assume the homogeneity of variances in the different groups).

### Survivability tests and electroretinogram (ERG)

For the survivability tests, 100 embryos per time point coming from a collective *srrm3* HET cross were grown in two tanks. At each time point all the fish were individually euthanized in 0.08% tricaine and placed into 100 uL NaOH 50mM for further DNA extraction as previously described. Then genotyping was performed in order to check the ratios of survivability of each genotype. For the darkness survivability test, 100 embryos coming from a collective *srrm3* HET cross were grown in 2 tanks covered with aluminum paper to mimic dark conditions (50 embryos per tank). At 13 dpf, the surviving fish were genotyped. The control experiment was performed growing embryos from the same *srrm3* HET cross in standard conditions.

For ERG experiments, *srrm3* MUT, *srrm4* MUT and DMUT larvae as well as different control siblings were recorded at 5 dpf as previously described ^72^. Briefly, larvae were dark adapted for at least 30 mins before recording. Then each of them was transferred to a piece of filter paper in a plastic recording chamber, which was filled with 1% agarose. The reference electrode was inserted into the agarose through a hole in the chamber. Afterwards, the eye ball of individual larvae was removed by a tungsten wire. The recording electrode, filled with E3 medium (5 mM NaCl, 0.17 mM KCl, 0.33 mM CaCl2, and 0.33 mM MgSO4) was placed against the center of the cornea. All these preparation steps were carried on under dim red light. The eye was then stimulated by 3 stimulation flashes. The duration of each flash was 100 ms and the interval between flashes was 12 seconds. The intensities were 0.6 lux, 6 lux and 60 lux (light source, HPX-2000, Ocean Optics). Electronic signals were amplified 1000 times by a pre-amplifier (P55 A.C. Preamplifier, Astro-Med. Inc, Grass Technology) with a band pass between 0.1 and 100 Hz and recorded via the self-developed National Instrument Labview program. The recordings were performed in 2 biological replicates for each genotype.

## Data availability

RNA-sequencing data was submitted to GEO under the project GSE180781. All other RNA-Seq samples used in this study are publicly available and listed in Table S13.

## Author contributions

L.C. designed the study under direct supervision of M.I., S.A.H. and L.S. F.M. contributed to concept and study design. L.C. performed the experiments and analyzed the data. L.C, S.A.H, F.M. and M.I. performed the splicing analysis. F.M. performed the evolutionary analysis, hypergeometric tests and GO analysis. L.L.B., C.R. and J.P. generated and characterized the mutant lines. C.R. handled zebrafish maintenance and performed the survivability tests. D.C. and S.A.H. performed the structural analysis. J.Z. performed the ERG and analyzed the data, under the supervision of S.N. S.C. performed the EM, analyzed the data and helped in evaluating and discussing results, under the supervision of S.Ba. S.J. helped in the apoptotic staining, under the supervision of V.R. S.Bo. performed experiments. S.M.V. did computational analyses. L.S. and M.I. provided resources. L.C. prepared figures and tables with the contribution of F.M. and S.A.H. L.C., S.A.H., F.M. and M.I. wrote the manuscript with inputs from all the authors.

## Acknowledgments

We thank Juan Valcarcel for the scientific support and critical reading of the manuscript; Xavier Hernandez-Alias and Antonio Torres-Mendez for the constant scientific discussion; Claire Lastrucci for supervision during the initial stages of the project; Xavier Henandez-Alias and Miquel Anglada-Girotto for assistance during the bioinformatic analysis; CRG Genomics Unit for the RNA sequencing services; Elena Polishchuk and the TIGEM Advanced Microscopy and Imaging Facility for the Electron Microscopy services. We acknowledge the support of the CERCA Programme/Generalitat de Catalunya and of the Spanish Ministry of Economy, Industry and Competitiveness (MEIC) to the EMBL partnership.

## SUPPLEMENTARY FIGURES

**Fig. S1 -.**
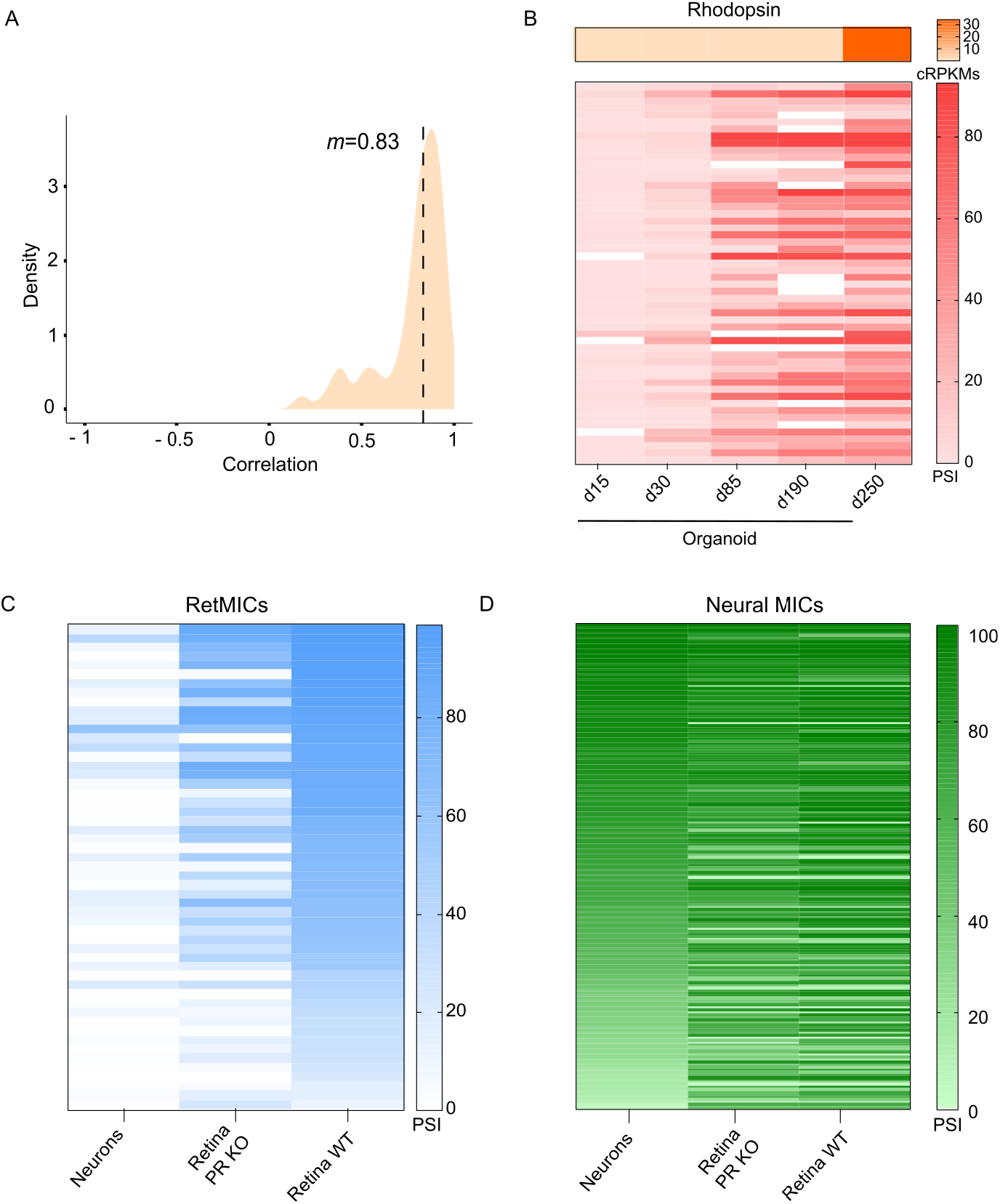
**(A)** Distribution of Spearman correlation coefficients between *Rhodopsin* expression and RetMIC inclusion levels (using the PSI metric) across the time course. Exons with insufficient read coverage in more than 40% of the samples were discarded; *m*, median of the correlations (n=54 valid exons). **(B)** Heatmap of *Rhodopsin* expression (top) and RetMIC inclusion levels (bottom) along organoid development (as described in Fig. 1C). RetMIC PSIs correspond to the average of three biological replicates for each time point. Values with insufficient read coverage (NA) are shown in white. **(C-D)** Heatmap of RetMICs (C) and and neural-enriched microexon (Neural MICs) (D) inclusion levels in hippocampal neurons, WT and *Aipl1* KO retinas. Events with insufficient read coverage were omitted.

**Fig. S2 -.**
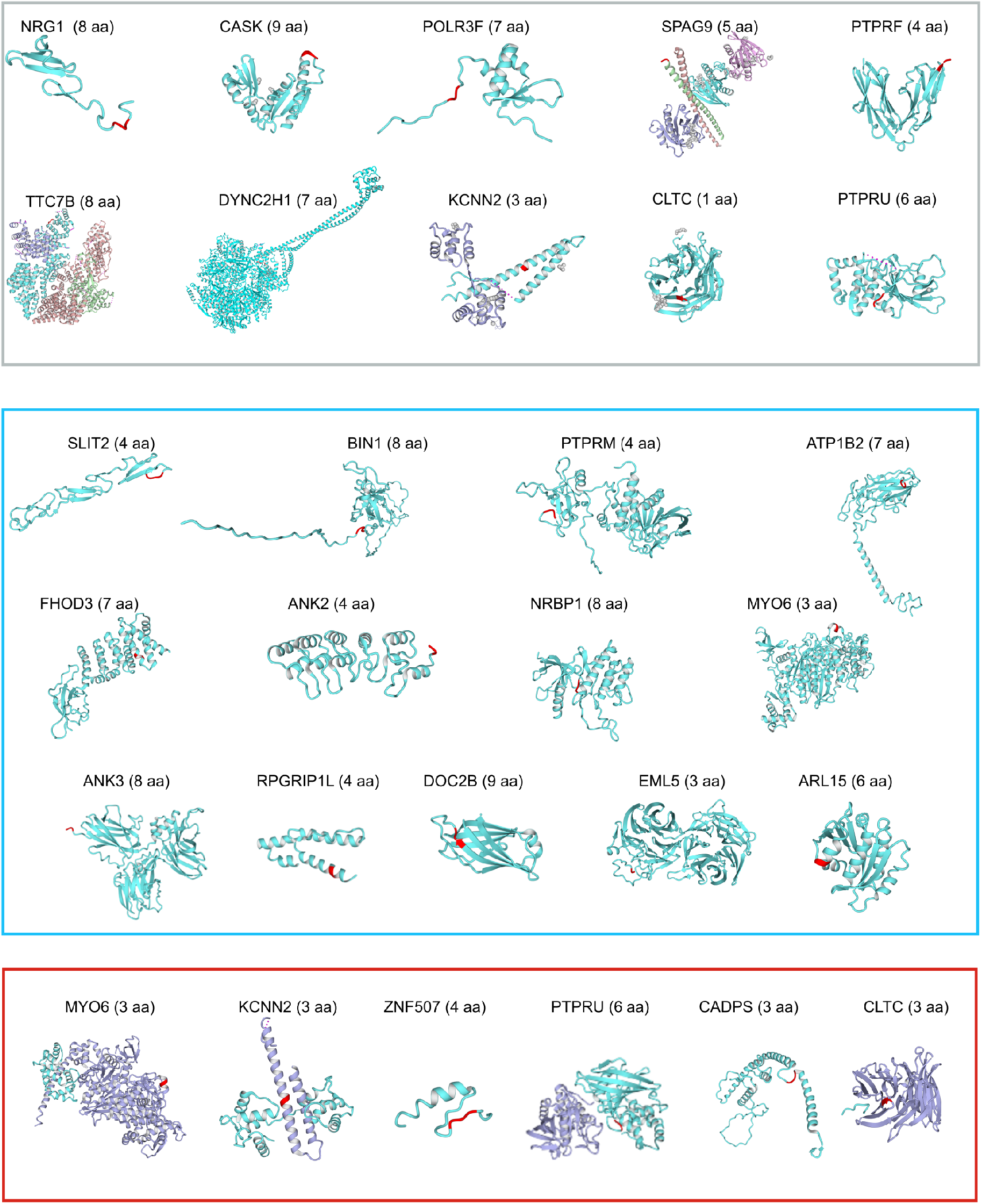
Protein 3D structures of RetMIC-containing genes for the exclusion isoform. Grey, blue and red boxes indicate experimental PDB structures, Interactome3D structures and ModBase models, respectively. The 3 residues around the RetMIC insertion sites are depicted in red. For each structure we indicate the gene name and the number of RetMIC-encoded amino acids (aa).

**Fig. S3 -.**
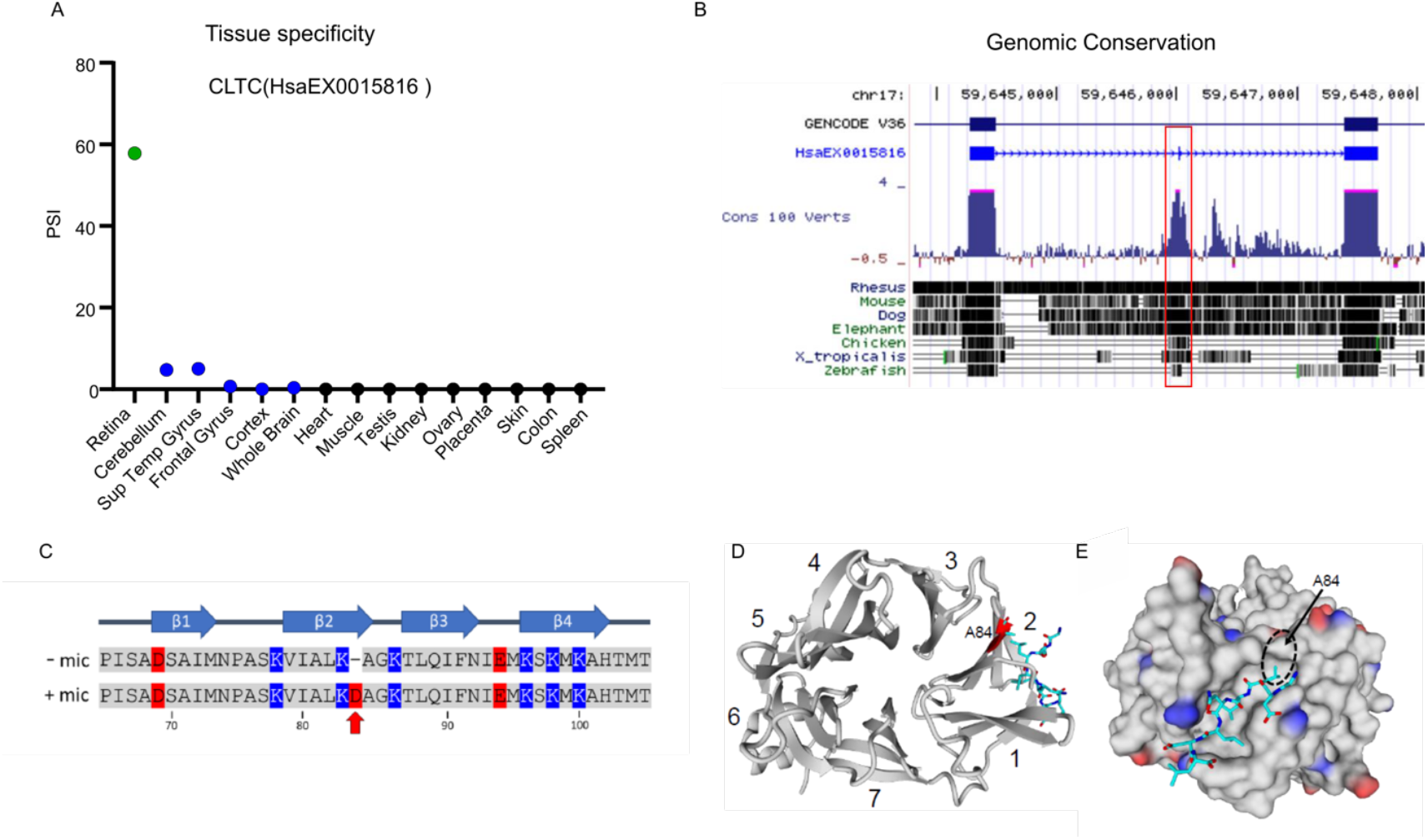
**(A)** Inclusion of a 3-nt microexon in *CLTC* (VastID: HsaEX0015816) across different tissues from *VastDB*. Retina is shown in green, neural tissues in blue. **(B)** Conservation of the microexon and neighboring intronic sequences in the genome across 100 vertebrate species, with seven representative species shown below. The flanking exons are shown as rectangles in the GENCODE V36 track, and the microexon is highlighted by a red box. Image generated from UCSC genome browser (http://genome.ucsc.edu). **(C)** Amino acid sequence alignment of *CLTC* with and without the microexon insertion, indicated by the red arrow. Negatively charged amino acids are depicted in red and positively charged in blue. The four beta strands comprising beta propeller blade #2 are indicated above the sequence. Amino acid numbering shown below corresponds to that of the *CLTC* gene without the microexon. **(D)** Ribbon structure of the CLTC terminal domain (PDB: 5M5R) with seven beta propeller blades numbered from N to C term. The AP2B1 clathrin-box peptide, which binds in the groove between blades 1 and 2, is represented by a stick model. The microexon-flanking amino acids of CLTC (K83 and A84) are highlighted in red, with the side-chain of A84 shown making hydrophobic contacts with the N-terminal lysine residue of the peptide clathrin box motif. Image generated by YASARA v19^73^. **(E)** Van der Waals surface of the CLTC terminal domain (PDB: 5M5R) showing the hydrophobic binding pocket of the AP2B1 clathrin-box peptide, with negatively charged side chains colored in red and positively charged side chains in blue. The hydrophobic side chain of A84 interacting with the N-terminal lysine residue of the peptide clathrin box motif of A2BP1 is circled. Insertion of the microexon results in a negatively charged Aspartic Acid residue at position 84, thereby disrupting this hydrophobic contact. Image generated by YASARA v19^73^.

**Fig. S4 -.**
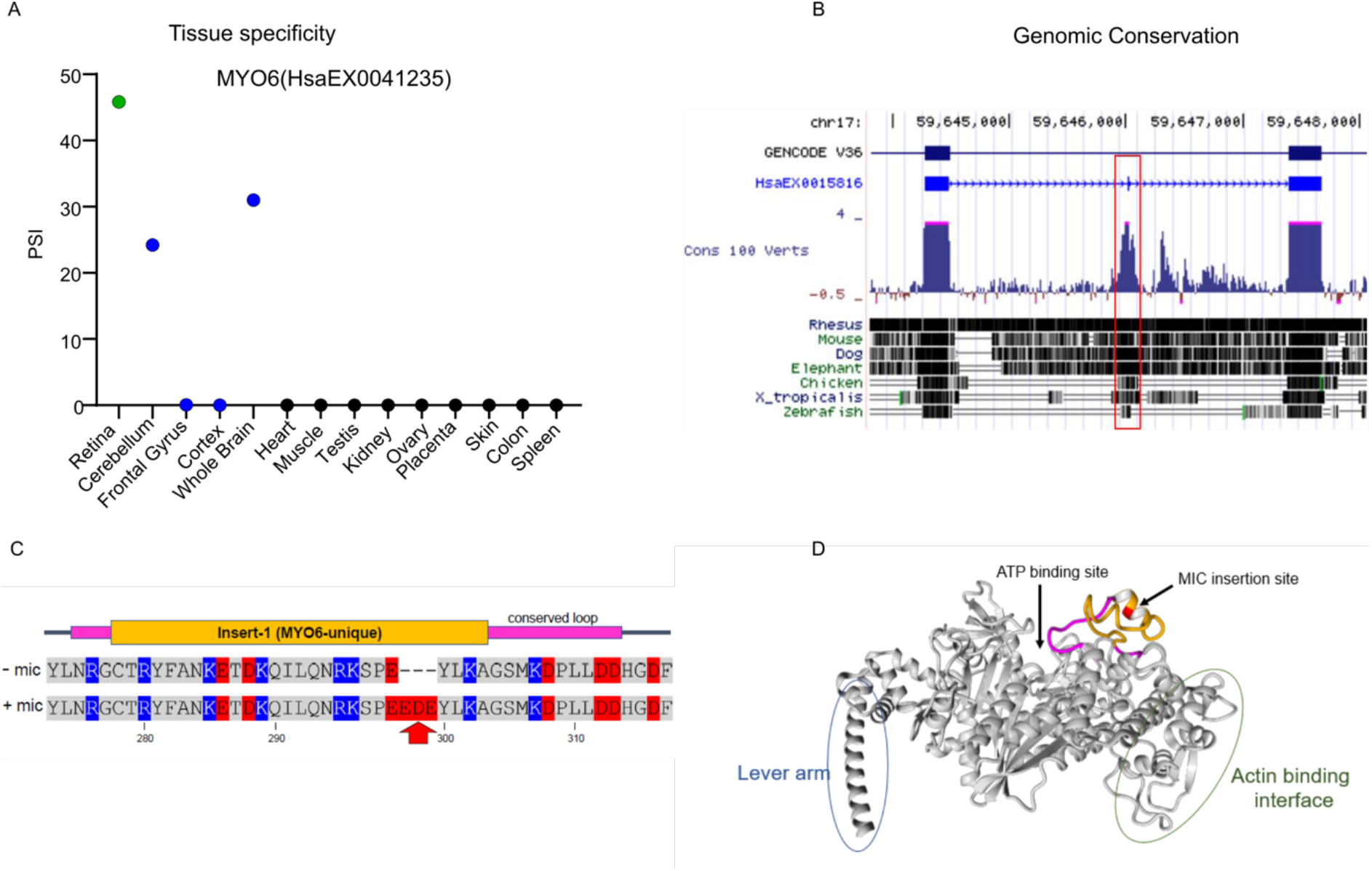
**(A)** Inclusion of a 9-nt microexon in MYO6 (VastID: HsaEX0041235) across different tissues from *VastDB*. Retina is shown in green, neural tissues in blue. **(B)** Conservation of the microexon sequence in the genome across 100 vertebrate species, with seven representative species shown below. The flanking exons are shown as rectangles in the GENCODE V36 track, and the microexon is highlighted by a red box. Image generated from UCSC genome browser (http://genome.ucsc.edu). **(C)** Amino acid sequence alignment of MYO6 with and without the microexon insertion, indicated by the red arrow. Negatively charged amino acids are depicted in red and positively charged in blue. The location of insert-1, which is unique to *MYO6* among other myosin genes, is indicated above the sequence in orange, while the remaining residues of the loop that are conserved among myosins are indicated in magenta. Amino acid numbering shown below corresponds to that of the MYO6 gene without the microexon. **(D)** Ribbon structure of the MYO6 catalytic motor domain (PDB: 2BKI), with insert 1 and the conserved loop colored as in Fig. S4D, and the microexon-flanking amino acids (E299 and Y300) highlighted in red. The close proximity of this loop to the ATP binding site is indicated by the black arrows. Image generated by YASARA v19^73^.

**Fig. S5 -.**
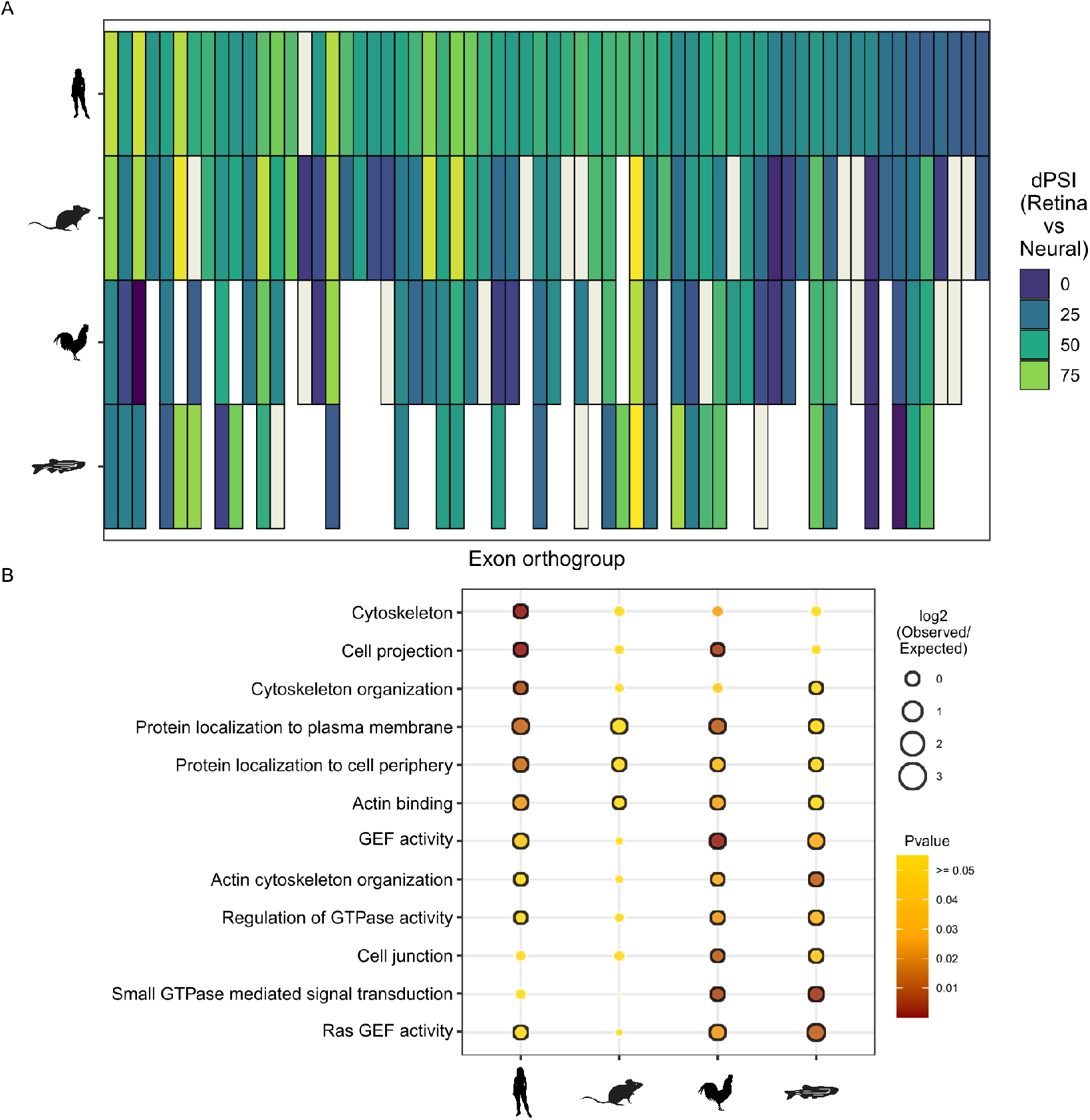
**(A)** Heatmap showing the retina-inclusion bias compared to neural inclusion of genomically conserved human RetMICs (top row) and their respective orthologs (i.e. exons belonging to the same exon orthogroup) in mouse, chicken and zebrafish. The color represents the delta PSI between the average of the retina samples and the average of neural samples considered for the definition of RetMICs. Human RetMICs and their selected orthologs are plotted in the same order as in Fig. 2C, to facilitate the comparison. Blanks and ivory rectangles indicate missing orthologs and missing ΔPSI values, respectively. **(B)** Dotplot representing functional enrichments for RetLONG genes across species. The functional enrichment of genes containing RetLONGs was separately tested for each of the species, and significant categories in at least two species (FDR-adjusted p-value ≤ 0.05) were plotted. The color reflects the adjusted p-value, with yellow color depicting p ≥ 0.05. The size of the dots is proportional to the log2 of the observed vs. expected ratio (O/E), and black borders around the categories highlight log2 O/E ≥ 1. GEF: Guanyl-nucleotide Exchange Factor.

**Fig. S6 -.**
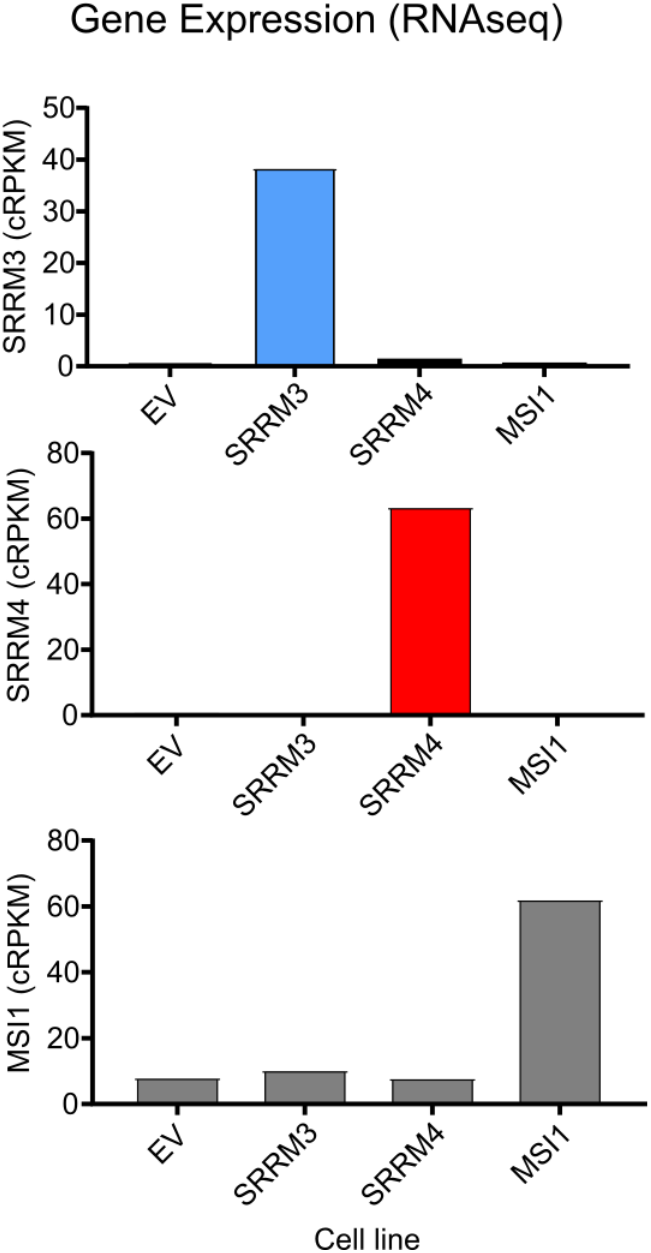
Normalized cRPKMs for *SRRM4, SRRM3* and *MSI1* in each cell line as measured from the RNA-seq data. EV: Empty Vector; cRPKMs have been quantile-normalized with the --NORM option in *vast-tools combine*.

**Fig. S7 -.**
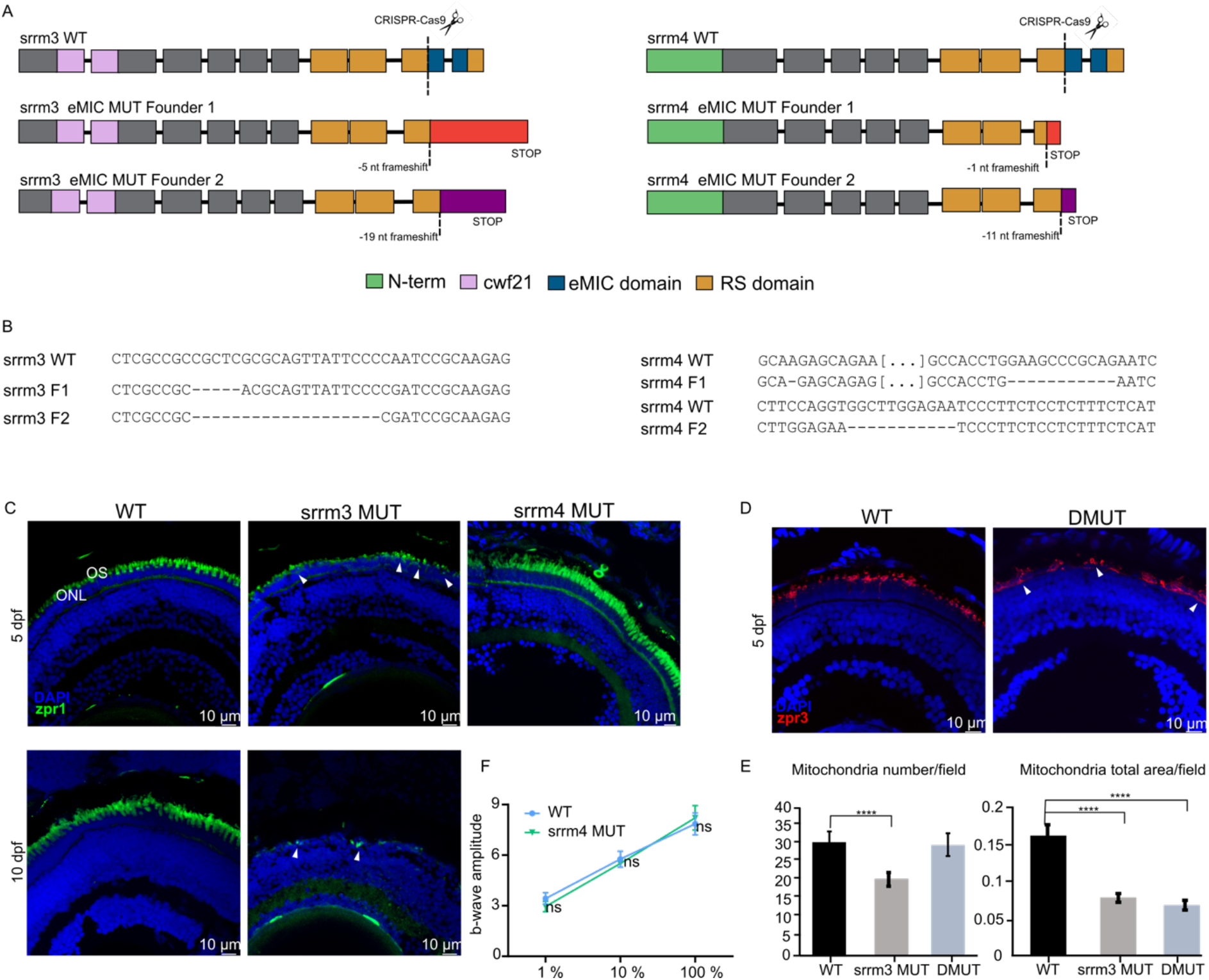
**(A)** Representation of the structure of *srrm4* and *srrm3* in the WT and MUT eMIC domain lines, highlighting the main protein domains; N-term: *srrm4* specific N-terminal domain; RS-domain: serine/arginine repetitive matrix domain; cwf21: PFAM domain binding to the spliceosome; eMIC: enhancer of microexons domain. **(B)** Sequences of *srrm3* and *srrm4* WTs and MUTs, highlighting the region of the mutation. For each gene, two founders have been generated. **(C)** ZPR-1 staining in *srrm3* MUT at 5 and 10 dpf. Arrows show Cone-opsin mislocalization. N = 6 for 5 dpf fish and N = 4 for 10 dpf fish. The right picture shows ZPR-1 staining for *srrm4* MUT at 5 dpf; N = 3. **(D)** ZPR-3 staining for DMUT at 5 dpf; N = 2. **(E)** Quantification of mitochondrial number/field and total mitochondria area/field in WT, *srrm3* MUT and DMUT. One-way ANOVA test with Tukey post-hoc analysis was applied (signif. codes: 0 ‘****’, 0.001 ‘***’, 0.01 ‘**’, 0.05 ‘*’). N ≥ 4 eyes/genotype. N ≥ 13 of fields/genotype was analyzed. **(F)** ERG recording from WT and *srrm4* MUT at 5 dpf, upon different light stimuli (1 %, 10% and 100%). Recordings were done in two biological replicates. N = 17 for WT and N = 23 for *srrm4* MUT.

**Fig. S8 -.**
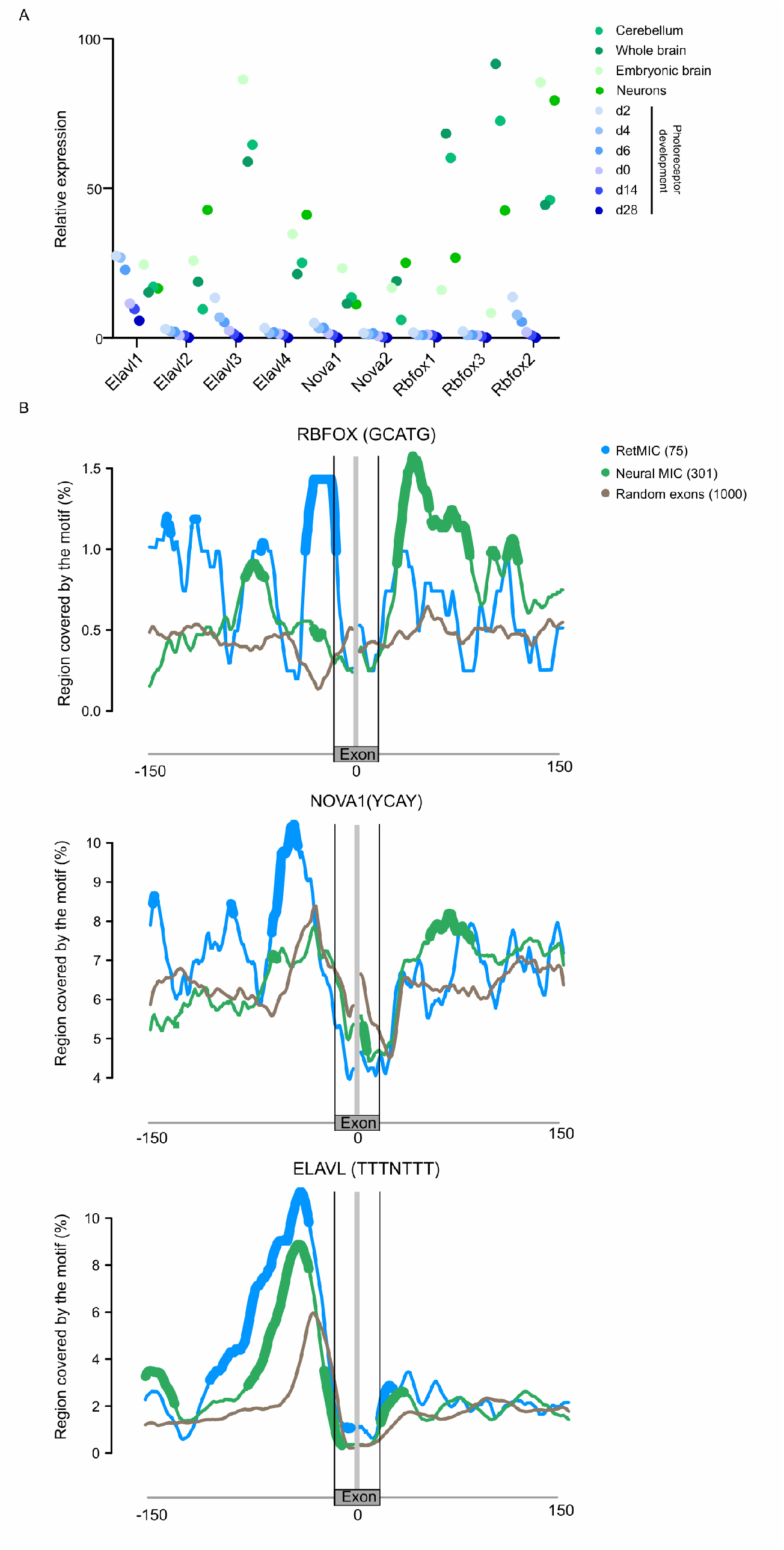
**(A)** Expression levels (cRPKMs) for *Elavl, Nova* and *Rbfox* family members across mouse developing rods, cerebellum, neurons, embryonic brain and whole brain samples (data from VastDB). **(B)** RNA maps of RBFOX, NOVA1 and ELAVL associated binding motifs in the regions surrounding RetMICs and neural-enriched microexons by length group and 1000 random exons. Regions with a significant difference in the motif coverage with respect to random (FDR< 0.05) are marked as thick lines.

