## Supplementary Materials for "Specialization of the photoreceptor transcriptome by *Srrm3*-dependent microexons is required for outer segment maintenance and vision"

**Supplementary Table legends**

**Table S1 - Information about vertebrate RetMICs and RetLONGs.** List of RetMICs and RetLONGs for all the studied species: human (hg38), mouse (mm10), chicken (galGal6) and zebrafish (danRer10). From columns 1 to 4, general information for each exon: gene name, event VastID, event genomic coordinate and exon length. Columns 5 to 16 contain sample and inclusion information: number of Retina replicates, number of other tissue groups (Others), average PSI in Retina, average PSI in Others, ΔPSI(Retina-Others), Standard Deviation (SD) of PSI distributions in Retina, SD of PSI distributions in Others, minimum average PSI in Others, maximum average PSI in Others, number of Neural samples, average PSI in Neural samples, SD of PSI distribution in Neural samples and the Retina Specificity Score (RSS). The last two sheets contain the inclusion of RetMICs and RetLONGs using the PSI metric in WT and *srrm3* MUT fish 5dpf eyes.

**Table S2 - GO enrichment analysis of genes containing RetMICs and RetLONGs.** Results of all GO enrichment analysis mentioned in the manuscript (enrichment of RetMICs and RetLONGs in human (hg38), mouse (mm10), chicken (galGal6) and zebrafish (danRer10); enrichment of Neural UP genes in human), performed through gprofiler2 in R as described in Methods. Each sheet contains: Genome assembly (Species), GO category name, Significance (TRUE if Pvalue $\leq 0.05$, FALSE otherwise), Pvalue (FDR corrected), Precision (Proportion of tested genes belonging to a given GO category), Observed (Number of tested genes belonging to a given GO category), Expected (Number of tested genes expected to belong to a given GO categories in a set of randomly chosen genes), Observed vs Expected (ratio Observed/Expected). GO categories are ordered by ascending Pvalue.

**Table S3 - Disease hypergeometric tests.** Settings, input values and results for the enrichment tests of the human RetMICs among genes associated with different retinal diseases.

**Table S4 - Domain annotations for RetMICs and RetLONGs.** VastDB automatic PFAM and PROSITE domain annotations for human RetMICs and RetLONGs. Annotations for all exons can be downloaded from VastDB. Events IDs are provided in column 1. C1 and C2 correspond to the exons upstream and downstream the alternative exon (A).

**Table S5 - Structural analysis of RetMIC-containing proteins.** 32 microexon events are listed with their relative gene name, Uniprot ID, predicted ORF disruption (yes/no). Structural model information as PDB identifier, title and molecule are given at columns 5-7. Modbase sub-models used in this study can be identified by their template structure and modelling quality respectively at column 5 and 6. Four proximal residues with relative solvent accessibility and secondary structure are listed at columns 8, 9 and 10. Column 11 indicates the upstream (C1) or downstream (C2) exons used to localise the target RetMIC within the structure.

**Table S6 - Regulation of retina-enriched exons by MSI1 and SRRM3/4 in cell lines.** Inclusion of RetMICs, RetLONGs (27nt < length ≤ 50nt) and RetLONGs (length >50 nt) among all the cell lines and experiments included in this study. Columns 1-4: name, event VastID, event genomic coordinate, exon length. Columns 5-14: for each comparison between the control and cells expressing MSI1 (red header), SRRM3 (purple) or SRRM4 (blue) ectopically, ΔPSI (PSI_OE - PSI_Cont) is provided. NA indicates the control and/or the experimental condition did not have sufficient read coverage to reliably estimate inclusion levels. Conditional formatting for ΔPSI values has been added from -100 (red) to 100 (green).

**Table S7 - Downregulated genes in srrm3 MUT eyes and associated enriched GO terms.** All downregulated genes in *srrm3* MUT eyes are listed, including Ensembl ID, gene symbol, expression levels (TPM) in WT and MUT siblings and log2 fold change (log2FC(MUT/WT)). The second sheet includes the GO enrichment analysis for the downregulated genes, considering only the genes with log2FC(MUT/WT) < -1.5 and performed using the software ClueGO. The sheet includes: GO ID, GO Term, Pvalue, Pvalue corrected with Bonferroni step down, GO Group, % of associated genes and number of genes. The different GO groups are highlighted in blue, green and grey as represented in Fig. 4C.

**Table S8 - Source data for the thickness and caspase analyses.** Outer Nuclear Layer (ONL) thickness measurements (µm) for *srrm3* MUT and WT retinas at 5 and 10 dpf are listed in the first sheet. For each fish included in the analysis, three values corresponding to the ONL length in three different areas of the retina are provided. The analysis of the Caspase3 staining for *srrm3* MUT and WT retinas is shown in the second sheet. For each fish, the number of apoptotic cells across the entire ONL are provided.

**Table S9 - Source data for the mitochondrial analysis.** EM analysis data for WT and mutant fish. The first sheet ("mitochondria area") includes the measurements of mitochondrial area for WT, *srrm3* MUT and DMUT fish. It also includes the data about the averaged mitochondrial area per field, the total mitochondria area per field and the number of mitochondria per field. The second sheet ("mitochondria perimeter") includes the measurements of mitochondrial perimeter for WT, *srrm3* MUT and DMUT fish, with the data about the averaged mitochondrial perimeter per field. The third sheet ("ipr space area") includes the measurements of interphotoreceptor (ips) space area for WT, *srrm3* MUT and DMUT fish.

**Table S10 - Source data for the ERG recordings.** ERG recordings data for WT and mutant fish. The first sheet ("srrm3 MUT") includes the recordings for WT, *srrm3* MUT (MUT) and heterozygous (HET) fish. The second sheet ("srrm3 srrm4 DMUT") includes the recordings from in-crosses of srrm4-/-;srrm3+/- fish. The labels (WT, MUT, HET) thus refer to the *srrm3* genotype with a *srrm4*-/- background. For each fish, three values are shown (1 %, 10 % and 100% light intensity). For each genotype, average, standard deviation (sd), total number of fish (n) and standard error of the mean (sem) are provided in the yellow boxes.

**Table S11 - Config file for calling tissue-enriched microexons.** Config files used to run *Get_Tissue_Specific_AS.pl* to derive sets of tissue-enriched microexons from PSI tables, for all the species considered in the manuscript. Column 3 "Excluded" contains the tissue groups (e.g. Retina, Muscle, Heart) excluded from the pool of compared tissues. This column must be empty to include all the tissues in the comparisons.

**Table S12 - Primers used in this study.**

**Table S13 - RNA-seq data used in this study.** For each species, SRA identifiers, read number, read length and source are provided for each public RNA-seq file used in this study. For RNA-seq samples generated in this study, mapping RNA statistics are provided in the last sheet.
